# Impact of 3-dimensional genome organization, guided by cohesin and CTCF looping, on sex-biased chromatin interactions and gene expression in mouse liver

**DOI:** 10.1101/577577

**Authors:** Bryan J. Matthews, David J. Waxman

## Abstract

**Background:** Sex differences in the transcriptome and epigenome are widespread in mouse liver and are associated with sex-bias in liver disease. Several thousand sex-differential distal enhancers have been identified; however, their links to sex-biased genes and the impact of any sex-differences in nuclear organization, DNA looping, and chromatin interactions are unknown.

**Results:** To address these issues, we first characterized 1,847 mouse liver genomic regions showing significant sex differential occupancy by cohesin and CTCF, two key 3D nuclear organizing factors. These sex-differential binding sites were largely distal to sex-biased genes, but rarely generated sex-differential TAD (topologically associating domain) or intra-TAD loop anchors. A substantial subset of the sex-biased cohesin-non-CTCF binding sites, but not the sex-biased cohesin-and-CTCF binding sites, overlapped sex-biased enhancers. Cohesin depletion reduced the expression of male-biased genes with distal, but not proximal, sex-biased enhancers by >10-fold, implicating cohesin in long-range enhancer interactions regulating sex-biased genes. Using circularized chromosome conformation capture-based sequencing (4C-seq), we showed that sex differences in distal sex-biased enhancer-promoter interactions are common. Sex-differential chromatin interactions involving sex-biased gene promoters, enhancers, and lncRNAs were associated with sex-biased binding of cohesin and/or CTCF. Furthermore, intra-TAD loops with sex-independent cohesin-and-CTCF anchors conferred sex specificity to chromatin interactions indirectly, by insulating sex-biased enhancer-promoter contacts and by bringing sex-biased genes into closer proximity to sex-biased enhancers.

**Conclusions:** These findings elucidate how 3-dimensional genome organization contributes to sex differences in gene expression in a non-reproductive tissue through both direct and indirect effects of cohesin and CTCF looping on distal enhancer interactions with sex-differentially expressed genes.

## Background

Sex differences in gene expression are found in several non-reproductive tissues, including the brain [1], immune system [2], kidney [3] and liver [4]. In the liver, sex differences in expression are associated with a higher incidence of aggressive liver cancer in males [5], increased susceptibility to autoimmune hepatitis in females [6], and sex differences in the metabolism of diverse pharmaceuticals and environmental chemicals [7]. Sex differences in the transcriptome and epigenome are best characterized in mouse liver [8, 9], where more than 1,000 genes [10], including many lncRNA genes [11, 12] and miRNAs [13], exhibit sex-biased expression regulated by the sex-differential temporal patterns of pituitary growth hormone secretion [9, 14]. Sex differences in the epigenome are widespread, and frequently are associated with sex differences in gene distal, but not gene proximal, regulatory elements, which show characteristic sex-differential patterns of histone marks and chromatin accessibility (DNase hypersensitive sites, DHS) [8, 15]. Three-dimensional looping is one mechanism that could potentially link the few thousand mostly distal sex-biased enhancers identified to individual sex-biased genes.

CCCTC-binding factor (CTCF) and the multi-protein complex cohesin are two major transcription factors regulating 3D genomic architecture. CTCF has primarily been studied for its role in DNA looping and insulation [16, 17], while cohesin is a molecular motor powering DNA looping via a loop extrusion mechanism [18, 19]. Loss of CTCF or cohesin is lethal in developing mouse embryos [20, 21]. However, when degradation of the cohesin loading factor Nipbl is induced in adult mouse liver, a dose-dependent loss of both cohesin binding and virtually all focal DNA looping is seen without major hepatocyte toxicity [22]. Loss of DNA looping also occurs in other systems following depletion of either cohesin [23] or CTCF [24]. Thus, CTCF and cohesin are both required for DNA looping.

The functional role of cohesin at a given genomic site is largely dependent on its binding partners. Cohesin lacks sequence-specific DNA binding activity, but is loaded and unloaded from chromatin by specific protein complexes [25]. Cohesin can participate in shorter-range looping between enhancers and promoters (‘gene loops’) in association with Mediator and tissue-specific transcriptional regulators [26-28]. Genomic sites bound by cohesin but not CTCF, i.e., cohesin-non-CTCF sites (CNC), tend to be highly tissue specific, but frequently show weaker binding than sites where cohesin and CTCF are both bound, i.e., cohesin-and-CTCF (CAC) sites [26, 29, 30]. Cohesin forms insulating loops at CAC sites, which have been variously characterized as Topologically Associating Domains (TADs) [31] and intra-TAD loops [32], loop domains [33] and insulated neighborhoods [27]. A majority of CAC-mediated insulated loops are conserved across tissues and cell types [31-34] and many are even conserved across mammalian species [35]. A distinct subset of CAC sites is important for enhancer-promoter interactions [36, 37], and may specifically help structure super-enhancers around key constituents enhancers, known as hubs [38]. The role of CTCF binding in the absence of cohesin, i.e., at Lone CTCF sites, is less clear. CTCF binds specific DNA motifs via its 11 zinc fingers [16], yet the function of Lone CTCF sites is likely also dependent on additional interacting proteins, such as the transcription factors YY1 [39], BORIS [40] and STAT5 [41].

CAC-mediated insulating loops are typically anchored by a pair of tightly bound CTCF sites oriented toward each other by the non-palindromic CTCF motif [33]. Supporting this finding, inversion or deletion of CAC-bound anchors results in a loss of looping [18, 42]. This stereotypical pattern of convergently-oriented CTCF binding enables the accurate prediction of CAC-mediated loops based on CTCF and cohesin binding activity and motif orientation alone [18, 32, 43]. Computational prediction of loops linking enhancers and promoters, i.e., gene loops, is a more elusive goal, although some recent progress has been made [44, 45]. Chromatin conformation capture technology using formaldehyde crosslinking and restriction enzyme digestion followed by proximity ligation can be employed to identify such loops experimentally, and thereby determine interaction frequencies between different genomic regions [46]. Circularized chromatin conformation capture with sequencing (4C-seq) is one such method that interrogates all potential interactions between a single site of interest and the rest of the genome [47].

Here we take a multi-pronged approach to elucidate the role of architectural proteins and 3D genome organization in regulating the widespread sex differences in gene expression seen in mouse liver. First, we identify sex-biased binding sites for cohesin and CTCF in mouse liver chromatin, a majority of which were found at intergenic sites distal to sex-biased genes. Further, we investigate the effects of a deficiency in cohesin loading in male mouse liver [22], and find that cohesin is specifically required for expression of male-biased genes with distal sex-biased regulatory elements. Finally, we use 4C-seq to directly evaluate sex differences in chromatin interactions involving sex-biased enhancers from five different genomic regions, and demonstrate the importance of loop domains for insulation of enhancer-promoter contacts at sex-biased genes. Overall, our findings highlight how 3-dimensional genome organization contributes to sex differences in liver gene expression in both direct and indirect ways.

## Results

### Sex differences in CTCF and cohesin binding in mouse liver

We used ChIP-seq to identify binding sites for CTCF and the cohesin complex protein Rad21 in both male and female mouse liver. We observed significant sex-difference in factor binding (>2-fold at FDR (false discovery rate) < 0.05) at 975 CTCF binding sites and at 1,011 cohesin binding sites (Fig. 1A; sites are listed in Table S1A and Table S1B). Peak regions showing significant sex differential binding for both CTCF and cohesin represent 13-14% of all 1,847 sex-differential sites (Fig. 1B; also see Fig. S1A for alternative filters to define sex-differential binding). An overall trend of sex-biased binding by both factors was seen for an even larger fraction of the sex-differential ChIP-seq peaks, as indicated by aggregate plots and heat maps (Fig. 1C).

**Fig. 1.**
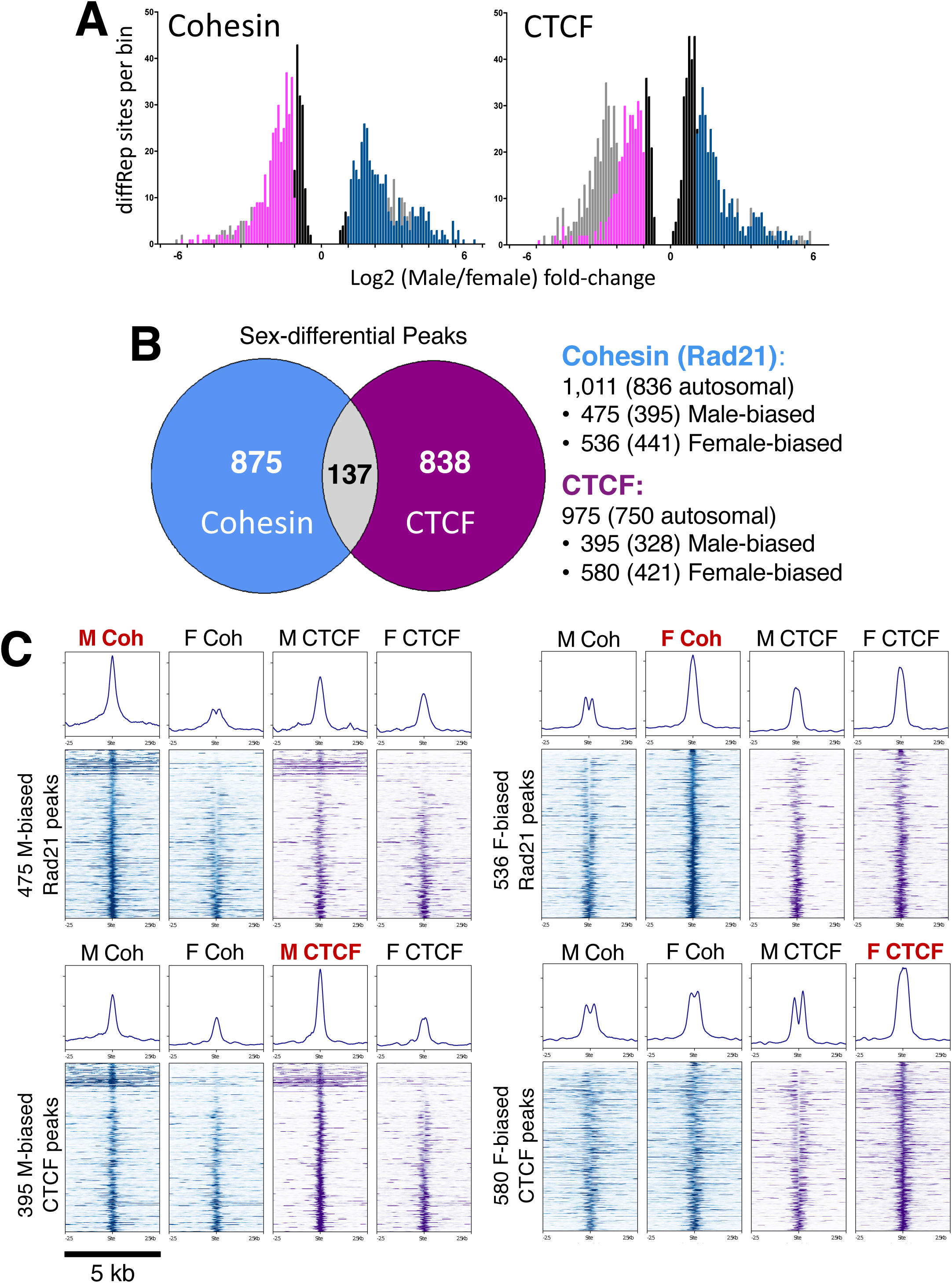
Sex differences in cohesin and CTCF binding to mouse liver chromatin. **A**. Distribution of male/female ratios for all diffReps-identified sex-differential sites that overlap a MACS2 peak for binding of cohesin (*left*) and CTCF (*right*). The y-axis shows the number of binding sites per bin, and the x-axis shows the sex difference in binding, expressed as log2(Male/Female) fold-change. Gray bars represent binding sites below the 10 read minimum count threshold, which were filtered out, and black bars represent sites that were statistically significant, but showed a |fold change| < 2 (values between −1 and 1 on the graph). Pink and blue bars respectively represent female-biased and male-biased sites above these thresholds. **B**. Venn diagram indicating 137 sex-differential peaks are common between cohesin and CTCF. Overlap is based on all sex-biased peaks, including male-biased and female-biased peaks on sex chromosomes (autosomal sex-biased peak numbers are shown in parenthesis, at the *right*). This pattern of limited overlap was also seen when the full set of unfiltered diffReps regions was examined (Fig. S1A). In total, 1,847 unique peaks exhibited significant sex bias in liver chromatin binding of CTCF and/or cohesin. **C**. Heat maps and aggregate plots for four sets of sex-biased cohesin (‘Coh’) or CTCF peaks. The peak set showing significant sex bias is highlighted in red at the top of each subpanel. For the heat map, read-in-peak normalized ChIP signals are shown for male and female cohesin binding (in blue) followed by male and female CTCF binding (in purple) within a 5 kb window centered around the differential peak summit. The aggregate profiles (*top*) represent the average signal of the heat map below for the same 5 kb window. Within each heat map, peaks are ranked based on the magnitude of sex bias from most sex-biased (*top*) to least sex-biased (*bottom*). Bimodal peaks, seen in the aggregate plots of several peak sets, is not a reflection of cohesin positioning relative to CTCF directionality, where the displacement is only ∼20 nt, as compared to the >100 nt shift observed here. Rather, it may be due to differences in co-binding of transcription factors in male or female liver displacing cohesin and/or CTCF within 1 nucleosome length.

Next, we classified the sex-differential CTCF and cohesin binding sites into the following groups: sites where both factors are bound (CAC sites) and that show sex-differential binding of either cohesin (ΔCoh) or CTCF (ΔCTCF), or both factors; cohesin-only binding sites (CNC sites) with sex-differential cohesin binding; and CTCF-only sites (Lone CTCF sites) with sex-differential CTCF binding. CAC sites comprised 45-66% of all sex-differential sites (Fig. S1B) and generally showed stronger factor binding than the sex-differential CNC and Lone CTCF sites (Fig. 2A, Fig. S2A). The strength of factor binding (Fig. 2A), the CTCF motif score (Fig. S2B), and the percentage of sites with a CTCF motif (Fig. S2C) were generally higher for the female-biased sites than the male-biased sites. In contrast, a higher fraction of male-biased than female-biased Lone CTCF sites contained a CTCF motif (66% vs 48%, Fig. S2C), but there was no significant sex difference in normalized ChIP signal or motif score (Fig. 2A, Fig. S2C). The latter sex differences may be driven by additional factors, such as the inhibitory effect of DNA methylation on CTCF binding [48, 49], where the same sequence motif in male and female liver could be preferentially-bound in males due to the hypermethylation of DNA seen in female compared to male mouse liver [50].

**Fig. 2.**
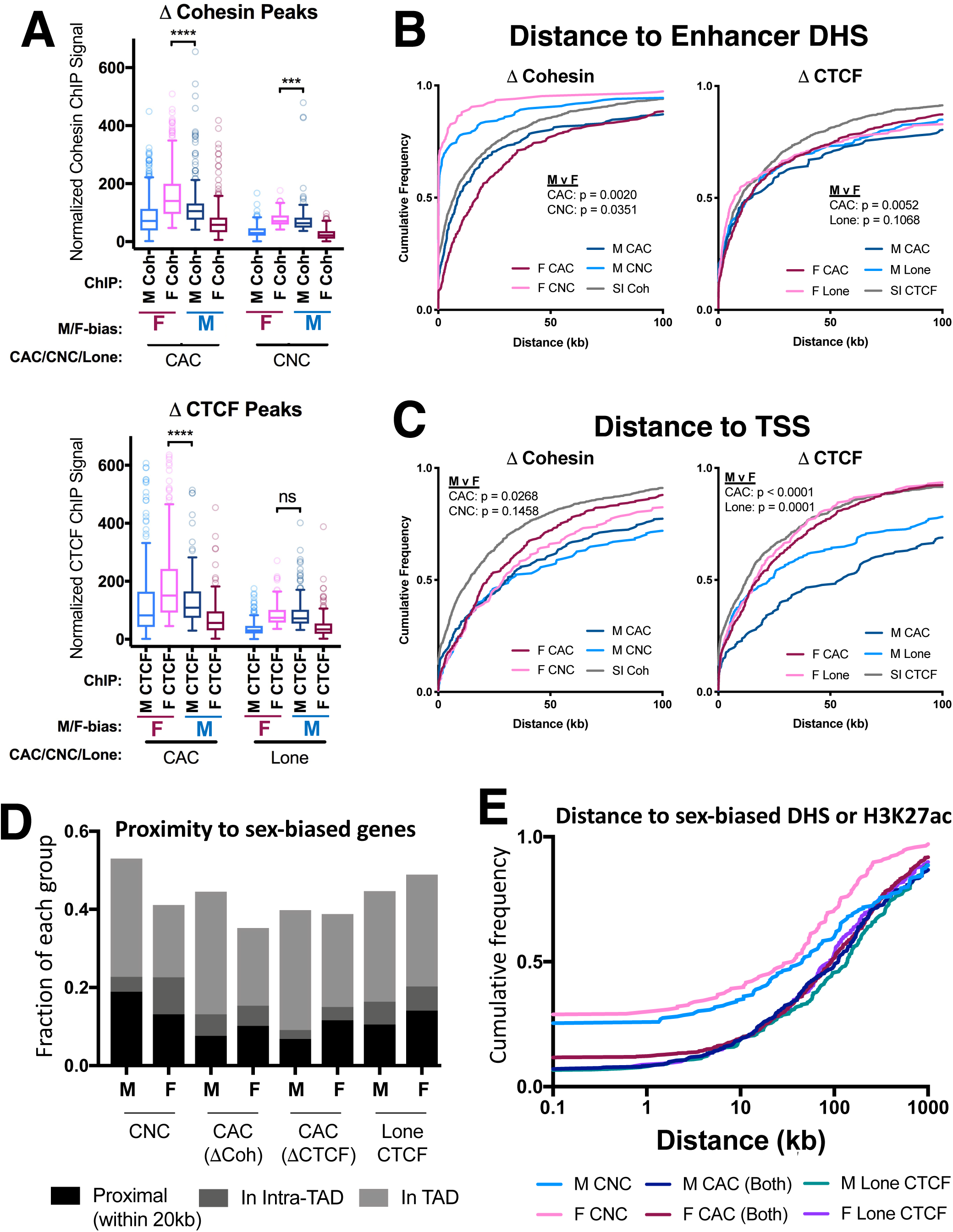
Cohesin and CTCF ChIP-seq binding strength and proximity to genes. **A**. Box plots of normalized ChIP-seq signal for the peak sets indicated on the x-axis. Peaks with sex differential binding for cohesin (*top graph*) and CTCF (b*ottom graph*) are shown. Each pair of boxplots represents the male and female ChIP-seq signal for the same set of peaks, defined by their sex bias and peak type (CAC or CNC, for ΔCohesin peaks; and CAC or Lone CTCF, for ΔCTCF peaks), as indicated below the x-axis. Peak scores were calculated by average intra-peak ChIP signal, normalized by total sequence reads per million in peak (RIPM; see Methods). Female-biased peaks were, on average, stronger than male-biased peaks by M-W test: p ≤ 0.001 for female vs male CAC(ΔCoh), CAC(ΔCTCF), and for CNC, but not for Lone CTCF peaks. **B**. Distance from each indicated set of cohesin and CTCF peaks to the nearest enhancer DHS. Cumulative frequency curves indicate the fraction of each group on the y-axis, within the distance in kb to the nearest enhancer DHS indicated on the x-axis. Enhancer DHS were defined based on their high ratio of the enhancer histone mark H3K4me1 over the promoter mark H3K4me3 at DHS [32]. Sex-biased CNC peaks are closer to enhancer DHS (median distance to eDHS of 0.22 kb for male-biased CNCs and 0.12 kb for female-biased CNCs; K-S pval < 0.0001 for all comparisons) than the other CTCF and cohesin peak classes (M CAC(ΔCTCF): 14.98 kb; F CAC(ΔCTCF) 13.76 kb; M Lone ΔCTCF: 13.88 kb; F Lone ΔCTCF: 7.17 kb). Female-biased CNC peaks are significantly closer to enhancer DHS than are male-biased CNC peaks (p=0.0351; K-S t-test). Male-biased CAC(ΔCohesin) peaks were closer to enhancers than female-biased CAC(ΔCohesin) peaks (p=0.002; K-S t-test), however, the reverse was found for CAC(ΔCTCF) peaks (p=0.0052; K-S t-test). Distance to nearest enhancer was not significantly different between male-biased and female-biased Lone CTCF peaks (p=0.1068; K-S t-test). P values for comparisons between male-biased and female-biased peaks of the same class are shown for each plot (K-S t-test). **C**. Distance from each indicated set of cohesin and CTCF peaks to the nearest TSS. Cumulative frequency curves indicate the fraction of each group on the y-axis within the distance in kb to the nearest TSS indicated on the x-axis. TSS for protein coding (RefSeq) and liver lncRNA genes were considered [70]. Female-biased cohesin and CTCF peaks are closer to TSS than male-biased CTCF and cohesin peaks of the same class (significance by K-S t-test is indicated at top left of each plot). Distance to the TSS was not significantly different for male-biased versus female-biased CNC peaks (p=0.1458; K-S t-test). **D**. Proximity of sex-biased cohesin and CTCF binding sites to sex-biased genes. Peak designations were as follows: Proximal, peaks < 20 kb from a sex biased gene TSS; Intra-TAD, peaks within the same intra-TAD loop as a sex-biased gene; or TAD, peaks in the same TAD as a sex-biased gene. Each of these groups is mutually exclusive. TAD loop [35] and intra-TAD loop [32] coordinates were from the indicated references. A set of 983 sex-biased biased protein-coding genes was used in this analysis (see Table S1 of [9]). **E**. Cumulative frequency curves show the fraction of each group (y-axis) within the distance in kb to the nearest sex-biased DHS or H3K27ac genomic region (x-axis), based on a merged list of published sex-biased DHS [15] and sex-biased H3K27ac ChIP-seq peaks [96] for male and female mouse liver. For this analysis, CAC peaks with sex-biased binding of CTCF and cohesin were combined and presented as a single group (CAC (Both)). Male-biased and female-biased CNC peaks are significantly closer to sex-biased DHS/H3K27ac than the four other peak classes (p < 0.001; K-S t-test). Female-biased CNC peaks were significantly closer to sex-biased DHS/H3K27ac than male-biased CNC peaks (p=0.0094; K-S).

### Mapping of sex-biased binding sites to TAD and intra-TAD boundaries, genes and regulatory elements

We investigated whether the sex-differential binding shown by cohesin and CTCF is linked to a sex-differential segmentation of the genome at the level of TAD and intra-TAD loops (DNA loop domains). We identified 137 CAC sites with significant sex differences in both CTCF and cohesin binding (Fig. 1B), of which 53 are on autosomes (Table S1C). 17 of the 137 sex-differential CAC sites overlap a TAD or intra-TAD loop anchor [32] in either male or female liver, of which 9 are on autosomes (Table S1C). Ten of the 17 CAC sites are associated with an intra-TAD loop predicted to be present in one sex only, of which 5 such loops contained one of more sex-biased genes. One of the 5 sex-based intra-TAD loop domains includes 6 sex-biased genes from the *Cyp2c* gene family and 2 sex-biased lncRNA genes (Fig. S2D). Consistent with the low frequency of sex-biased CAC sites at TAD or intra-TAD loop anchors, 88-93% of all intra-TAD loops and anchors predicted in male liver were also predicted in female liver (Fig. S2E, Table S1J). We conclude that a large majority of CAC-mediated intra-TAD loops are conserved between the sexes in mouse liver, and thus, sex differential intra-TAD loop domains do not comprise a major regulatory mechanism for sex-biased gene expression. Nevertheless, TAD and intra-TAD loop domains that are common to both sexes may indirectly guide enhancer-promoter looping in a sex-biased manner, as described below.

Next, we considered whether the sex-biased binding sites for CTCF and cohesin might help link sex-biased regulatory elements to sex-biased gene promoters. We examined the locations of these sites relative to enhancer DHS, defined as open chromatin regions (i.e., DHS) with a high ratio of H3K4me1 to H3K4me3 ChIP-seq signal [32]. We found that sex-biased CNC sites are much closer to enhancer DHS (median distance of 202 bp (in males) or 119 bp (in females)) than were sex-biased CAC sites (median distance = 8.8 kb and 17.5 kb, for male-biased and female-biased CAC, respectively) (Fig. 2B, *left*). Thus, although sex-biased CNC sites are weaker binding than CAC sites (Fig. 2A), a majority are found at enhancers and may be functional. Female-biased CTCF binding sites tended to be closer to the transcription start site (TSS) than male-biased sites (Fig. 2C), despite similar ChIP-seq sample quality (Table S3B).

All classes of sex-biased cohesin and CTCF binding sites were primarily distal from sex-biased genes: < 20% mapped within 20 kb of a sex-biased gene, and only 35 to 53% were found within the same TAD and could therefore be considered potential *cis* regulators of sex-biased genes (Fig. 2D). Consistent with the association of cohesin with enhancers [26, 51], 25-29% of sex-biased CNC sites overlapped a sex-biased enhancer (either a sex-biased DHS or a sex-biased H3K27ac peak), as compared to only 7-11% of sex-biased CAC and Lone CTCF sites (Fig. 2E). Furthermore, 77% (72 of 96) of sex-biased CNCs that overlap a sex-biased enhancer are >20 kb from a TSS of sex-biased gene (median distance, 238 kb). However, a majority of all classes of sex-biased cohesin and CTCF sites are quite distant from sex-biased regulatory elements (Fig. 2E), consistent with their being distal regulators of sex-biased gene expression.

Sex-biased liver CNC sites, as well as Lone CTCF sites, showed much more tissue-specific binding of CTCF across mouse tissues [52, 53] than did sex-biased liver CAC sites (Fig. S3A, lower vs upper panels). In addition, liver-expressed genes that mapped to sex-biased CNC sites showed a more liver-specific expression pattern than genes mapping to sex-biased CAC sites, or liver-expressed genes overall (Fig. S3B). This suggests that sex-biased CNC sites participate in tissue-specific transcriptional regulation, as was described for CNC peaks generally [26, 29, 30]. Significant differences in the tissue-specificities of CTCF binding were also seen at male-biased compared to female-biased CAC sites and Lone CTCF sites (Fig. S3A, Fig. S3C).

### Impact of cohesin depletion on distally-regulated male-biased genes

As noted above, 35-53% of sex-biased cohesin and CTCF binding sites are within the same TAD as at least one sex-biased gene (Fig. 2D) and could play a role in DNA looping between sex-biased enhancers and sex-biased gene promoters. Examples of sex-biased CTCF and/or cohesin binding sites that were either proximal (< 20 kb) or distal to sex-biased genes are shown in Fig. 3A-3C. Fig. 3A shows a highly female-biased enhancer (female-biased DHS and female-biased H3K27ac marks) with an overlapping female-biased CAC site (green arrows) located 33 kb upstream of the female-biased gene *Slc22a29* (F/M expression ratio=8.7), the closest TSS. Fig. 3B shows *Cml5*, a male-specific gene (M/F expression ratio = 20.2) with a male-biased CNC site that overlaps a male-biased DHS ∼3 kb upstream of its TSS (Fig. 3A, red arrow). The adjacent gene, *Nat8* (M/F = 4.2), has a male-biased CAC site that overlaps a male-biased DHS located ∼12 kb upstream of the TSS (Fig. 3B, green arrow). Conceivably, the sex-biased binding of cohesin and CTCF at these sites could contribute to looping of the associated sex-biased enhancers to their correspondingly sex-biased gene targets. Finally, Fig. 3C shows two male-biased complement C8 genes (*C8a* (M/F = 3.3) and *C8b* (M/F = 2.8)) that are linearly quite distant (∼1.5 Mb) from a cluster of strongly male-biased enhancers near the 5’ end of the same TAD. The TAD structure of this genomic region suggests these enhancers are spatially more proximal to the *C8* genes than they are to than the linearly much closer *Oma1* gene, located just inside the adjacent TAD (also see Fig. S4A).

**Fig. 3.**
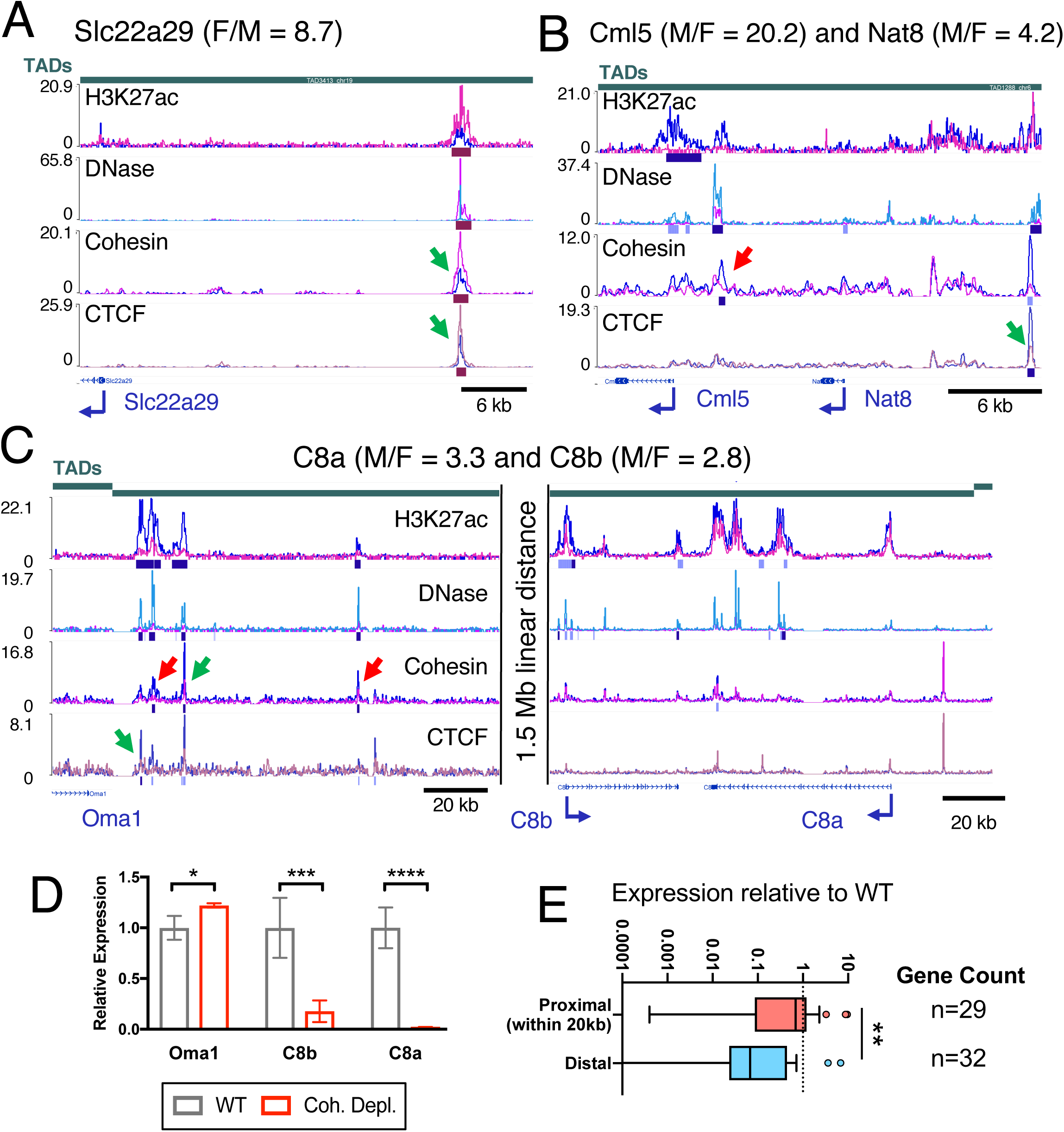
Proximal and distal sex-biased regulatory elements and impact of cohesin depletion. Shown are WashU Epigenome Browser screenshots of sex-biased cohesin and CTCF binding events proximal (≤ 20 kb from a sex-biased gene TSS) and distal (> 20 kb) to sex-biased genes in mouse liver. Tracks shown are (from *top* to *bottom*): TAD and TAD boundary location [indicated by a vertical shift in the TAD track, as on the *left* and on the *right* sides of panel C], H3K27ac ChIP-seq, DNase-seq, cohesin (Rad21) ChIP-seq, CTCF ChIP-seq, and Ref-seq genes. ChIP-seq and DNase-seq data in each track is shown superimposed for male (blue) and female (pink) liver after normalization to total reads per million in the union of peaks for that factor (see Methods). Below each ChIP-seq track, a horizontal bar identifies genomic regions that show significant male-bias (blue bar) or female-bias (pink bar), with darker and lighter shades indicating strict and lenient cutoffs for sex-bias, respectively (see Methods). Green arrows indicate CTCF sex-differential CAC, and red arrows indicate CTCF sex-differential CNC and Lone peaks. Male/Female stranded polyA+ RNA-seq gene expression ratios [11] are indicated above each panel. **A**. *Slc22a29* is a female-biased gene with a distal female-biased non-anchor CAC peak overlapping a robust female-biased DHS with female-biased H3K27ac histone mark accumulation ∼34 kb upstream (green arrow). The region shown spans chr19:8290981-8333632. **B**. Two male biased genes, *Cml5* and *Nat8*, with proximal male-biased cohesin and CTCF peaks overlapping male-biased DHS (red, green arrows). *Cml5* has a ∼3 kb upstream male-biased CNC peak overlapping a strongly male-biased DHS. The *Cml5* promoter shows strong male-biased H3K27ac marks and a weaker male-biased DHS. *Nat8* has a weakly male-biased DHS at its promoter and a stronger male-biased DHS ∼12 kb upstream that overlaps a male-biased CAC peak. The region shown spans chr6:85766132-85794443. **C**. Male-biased genes *C8a* and *C8b* have distal male-biased CAC and CNC peaks overlapping male-biased DHS and H3K27ac marks (green and red arrows, respectively). The linear distance between the upstream male-biased CAC peaks and downstream male-biased genes is > 1.5 Mb. *C8a* and *C8b* reside on opposite ends of the same TAD. *Oma1* is close in linear distance to these sex-biased regulatory elements, but shows no sex differences in expression; based on TAD structure it is not be predicted to interact with the highlighted male-biased enhancers. See Fig. S4A for the full length of the TAD and a model of spatial positions. The left portion of this figure spans chr4:103027454-103167067, and the right portion spans chr4:104433514-104583344. **D**. Loss of cohesin binding decreases expression of *C8a* and *C8b* significantly (p ≤ 0.0001 and p=0.0020; M-W t-test), while expression of *Oma1* increases (p=0.0102; M-W t-test). Bars represent the mean expression of the group for a given gene relative to the mean of the WT group (equal to 1), and error bars show the standard deviation based on n=4 per group. **E**. Loss of cohesin binding has a 10-fold greater suppressive effect on male-biased genes with distal sex-biased enhancers than those with proximal sex-biased enhancers. Shown is the mean expression for cohesin-depleted versus wild-type liver, such that a value of 0.1 represents a 10-fold reduction in expression after cohesin loss. The median relative expression for DHS/H3K27ac-proximal genes is 0.69 (representing a modest suppressive effect of cohesin loss) and the corresponding median for DHS/H3K27ac-distal genes is 0.07, indicating a >10-fold greater reduction in gene expression (p = 0.0087; M-W). Similar results were obtained when the definition of proximally-regulated genes was relaxed to include genes with a TSS < 20 kb from *<u>either</u>* a male biased DHS or a male biased DHS (median of 0.45 versus 0.042 for distal genes). Also see Table S2.

Loss of cohesin binding in male mouse liver, achieved by depletion of the cohesin loading factor, Nipbl, leads to a loss of distal enhancer-promoter contacts and an increase in local ectopic contacts, which can activate proximal genes [22]. We compared the effects of cohesin loss on the expression of male-biased genes with proximal sex-biased enhancers versus those that have only distal (>20 kb) sex-biased enhancers. Fig. 3D shows the relative changes in gene expression in cohesin-depleted compared to wild-type male mouse liver for the three genes included in Fig. 3C. Expression of *C8a* and *C8b* decreased, by 98% and 82%, respectively, upon loss of chromatin-bound cohesin in male liver, while expression of *Oma1* increased modestly (+22%), perhaps by an enhancer hijacking mechanism [54]. In contrast, the two male-biased genes with proximal sex-biased enhancers, *Cml5* and *Nat8* (Fig. 3A), showed no significant change in expression following cohesin loss (Fig. S4B).

Next, we examined a set of 61 male-biased genes and verified the requirement of cohesin for expression of the distally-regulated but not the proximally-regulated male-biased genes: male-biased genes with distal (> 20 kb) sex-biased regulatory elements were significantly more sensitive to loss of cohesin than male-biased genes with proximal sex-biased enhancers (median decrease in expression upon cohesin loss: 14.3-fold vs. 1.4-fold; Fig. 3E). This finding likely results from a requirement for cohesin for distal interactions, via either a direct or an indirect looping mechanism. Conceivably, for sex-biased genes with nearby sex-biased regulatory elements, enhancer-promoter loops required for gene expression can be maintained over short genomic distances by transcription factors such as Mediator [26] or YY1 [55], and without a need for cohesin.

### 4C-seq analysis of sex-biased chromatin interactions in mouse liver

Next, we implemented 4C-seq analysis centered on six viewpoints at sex-biased enhancers in five distinct genomic regions to determine whether a sex-bias in enhancer−promoter loops is associated with sex-biased liver gene expression. Our findings (Fig. 4) are based on n=3 individual biological replicates per sex, whose 4C-seq interaction profiles are displayed in Fig. S5.

**Fig. 4.**
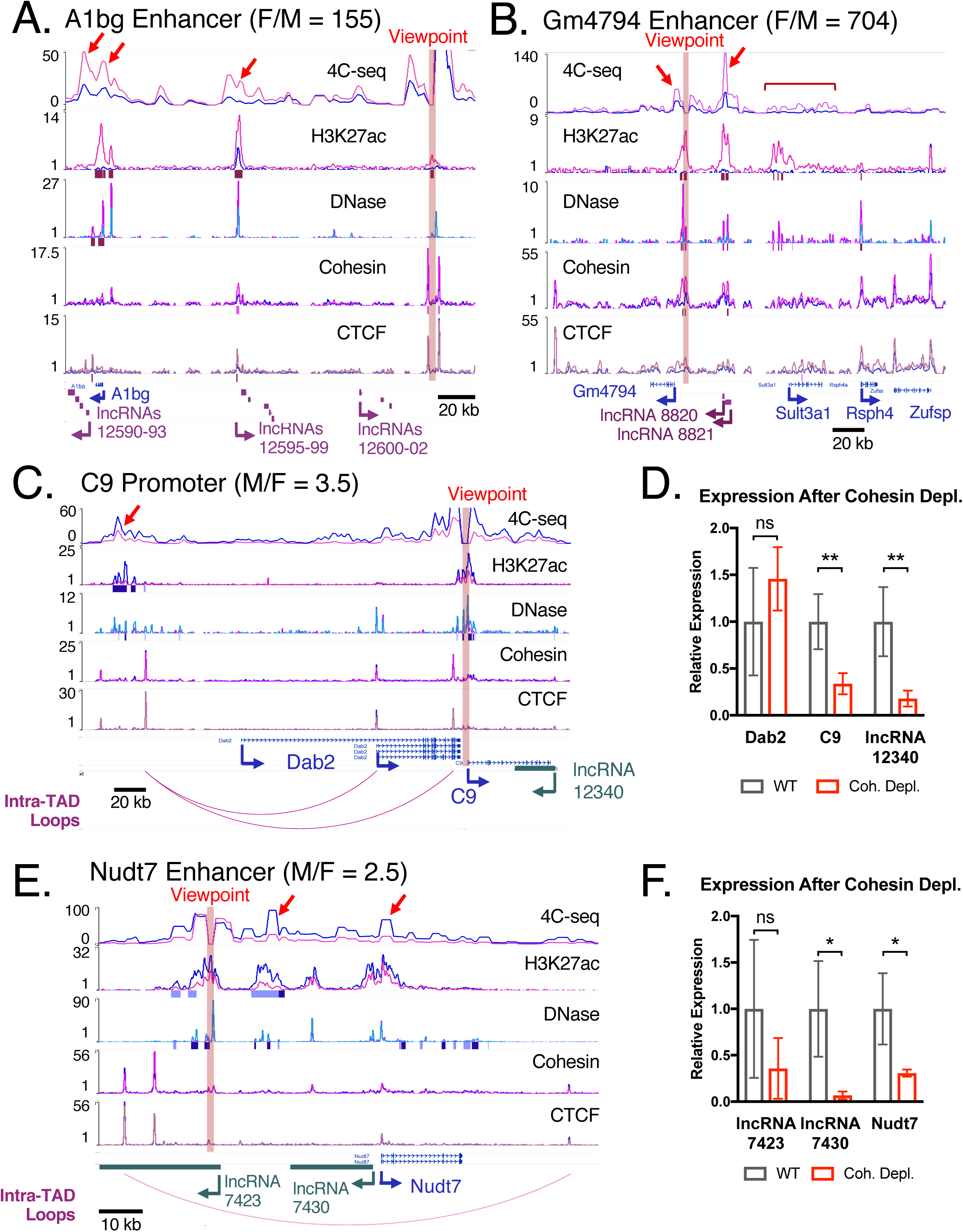
4C-seq analysis of two female-biased and two male-biased genes. A, B, C, E. Shown are WashU Epigenome Browser screenshots for four genomic regions investigated by 4C-seq. The *upper track* presents 4C-seq data for four viewpoints, marked by a vertical highlight in each panel. The 4C-seq track is based on merged data from three biological replicates for each sex, calculated from the median value for a sliding window of 11 restriction fragments, in reads per million normalized 4C-seq signal per sex. These values are overlaid for visualization using the Matplot functionality built into the WashU genome browser with default parameters. The next four tracks show normalized DNase-seq or ChIP-seq signal for the indicated factors, and correspond to those described in Fig. 3. Sex-biased lncRNAs are shown below the Refseq gene track in pink (female-specific lncRNAs) or blue (male-specific lncRNAs). C and E also show locations of intra-TAD loops (pink arcs) below the Refseq gene track. 4C-seq data for individual biological replicates is presented in Fig. S5. **A**. Distal enhancer viewpoint near *A1bg* and female-biased lncRNAs. The region shown includes 12 female-biased and nuclear-enriched mono-exonic lncRNAs, which fall into three clusters. The lncRNAs in each cluster are all transcribed from the same strand, as indicated by the arrow marking the TSS and direction of transcription of the most upstream lncRNA in each cluster. LncRNAs 12590-12593 show the strongest 4C-seq interactions with the viewpoint and also the most consistent female bias (Fig. S5E, Fig. S5F). The region shown spans chr15:60733512-60954051. **B**. Enhancer viewpoint 8 kb upstream of *Gm4794* interacts primarily with the proximal promoter (*left* red arrow) and a strong female-biased enhancer (*right* red arrow). Viewpoint enhancer and the interacting enhancer both contain female-biased CNC peaks (marked below cohesin track in dark pink). *Sult3a1* shows a weak and broad pattern of interaction with this viewpoint (red bracket). The region shown spans chr10:33418446-33680888. **C**. Viewpoint at the promoter of *C9* interacts with a distal male-biased enhancer, bypassing the gene *Dab2*. This viewpoint shows a strong male-biased interaction with distal male-biased enhancer indicated by a red arrow. The nested intra-TAD loop structure (loops at bottom) may facilitate the high and male-biased expression of *C9* by looping out the intervening gene *Dab2*. There is a weaker, but apparent male-biased interaction with the TSS of the short isoform of *Dab2*. Based on CAGE data and Refseq annotation, the shorter isoform of *Dab2* is predominantly expressed in liver (data not shown). The region shown spans chr15:6147917-6461799. **D**. Cohesin depletion significantly reduces expression of *C9* and its antisense lncRNA (*lnc12340*): 66% reduction, p=0.0056; and 82% reduction, p=0.0049, respectively. Bars represent mean expression compared to wild-type liver (WT, set equal to 1), with error bars showing standard deviation for n=4 per group. **E**. *Nudt7* lacks male-biased DHS or H3K27ac at its promoter, but interacts with a distal male-biased enhancer within the same intra-TAD loop. This viewpoint is anchored at a male-biased DHS (red highlighting). Prominent interactions with this viewpoint (red arrows) are with a neighboring male-biased enhancer and the promoter of *Nudt7*. While the TSS of *lnc7430* and *Nudt7* are close, the viewpoint enhancer interacts specifically with the TSS of *Nudt7*. The region shown spans chr8:116592444-116707613. **F**. Upon loss of chromatin-bound cohesin, expression of both *Nudt7* and its bi-directionally transcribed lncRNA (*lnc8430*) are significantly reduced (69% decrease, p=0.0116; and 93% decrease, p=0.0113, respectively). Expression of *lnc7423* was not significantly impacted. Data presentation is as described in D.

#### *A1bg* region

We anchored a 4C-seq viewpoint at a female-biased enhancer near the strongly female-biased gene *A1bg* (F/M = 155) (Fig. 4A, vertical pink highlight; Fig. S5A). There are 12 female-biased, mono-exonic nuclear-enriched lncRNAs [12] in three clusters across the genomic region displayed. Robust interactions were observed in female but not male livers between the viewpoint enhancer and three genomic regions (red arrows): a strong female-biased enhancer (*right* arrow), the promoter of *A1bg* (*middle* arrow), and a region downstream of *A1bg* that contains a cluster of four female-specific lncRNAs (lncRNAs 12590-12593; *left* arrow), where we observed the strongest interactions. The lncRNAs in this cluster are more highly expressed (Fig. S5E) and are more consistently female-biased across various RNA-seq datasets than the other two lncRNA clusters (Fig. S5F). The maximum expression of these 12 lncRNAs ranged from 0.31 to 2.71 fragment per kilobase per million sequence reads (FPKM) in female liver compared to 0 to 0.02 FPKM in male liver (Fig. S5E).

The precise relationship between the female-biased expression of these lncRNAs and the female-bias in 3D interactions with the distal enhancer is not known. The interaction may be regulatory in nature (e.g., an enhancer-promoter interaction, as with any gene) or it could be facilitated by one or more of the 12 nuclear-enriched, female-biased lncRNAs, as was described for other lncRNAs (*Xist* [56], *Firre* [57], *Haunt* [58]). Alternatively, the female-specific interactions shown may be primarily those of regulatory enhancers driving expression of female-specific genes, including *A1bg* and multiple lncRNAs. The female-biased CTCF binding seen at both interacting regions (*right* and *left* arrows) lends mechanistic support to the latter proposal, with CTCF mediating enhancer-promoter and enhancer-enhancer interactions. As CTCF can interact with lncRNAs in a functional manner, and with high affinity [59], these two mechanisms are not mutually exclusive; one or more of these highly female-specific lncRNAs (Fig. S5E) could function in a *cis*-acting manner to selectively guide CTCF binding and interactions unique to female liver.

#### Sult3a region

We used 4C-seq to interrogate an enhancer viewpoint proximal to the highly female-specific gene *Gm4794* (F/M = 704; also known as *Sult3a2*) and also two female-specific mono-exonic lncRNAs, *lnc8820* and *lnc8821* (F/M = 34 and 90, respectively) (Fig. 4B, Fig. S5B). The enhancer viewpoint is distal to two other female-specific protein coding genes, *Sult3a1* (F/M = 288) and *Rsph4a* (F/M = 34). We observed female-biased 4C-seq interactions with the proximal promoter of *Gm4794* (*left* red arrow) and with strong female-biased enhancers that overlap *lnc8820* and *lnc8821*. Unlike the enhancer nearby *A1bg*, these interactions are associated with female-biased CNC peaks present at both the viewpoint enhancer and the downstream interacting enhancer. Despite the absence of CAC insulator elements across this genomic region (and consequently, the absence of intra-TAD loops), the observed focal interactions are all local, within ∼35 kb of the viewpoint. However, the viewpoint enhancer also made weak interactions to a broad, ∼45 kb region extending from ∼15 kb upstream of the promoter of *Sult3a1* to ∼10 kb beyond the gene body (Fig. 4B, red bracket). This 45-kb region contains several robust female-biased DHS and H3K27ac peaks, but lacks cohesin binding, which may account for the lack of strong, focal interactions with the viewpoint enhancer. *Gm4794* and *Sult3a1* may both interact with the enhancer that overlaps *lnc8820* and *lnc8821*, but this weak (and perhaps indirect) association cannot be captured by proximity ligation under standard 4C-seq conditions. Reciprocal 4C-seq experiments anchored at these lncRNA TSS or alternative 4C methods with increased sensitivity [60] may be needed to validate these weaker interactions.

#### *C9* region

We examined a 4C-seq promoter viewpoint placed at the complement factor *C9* gene (M/F = 3.5) to investigate whether intra-TAD loops can indirectly coordinate enhancer-promoter contacts preferential to one sex in a genomic region devoid of sex-biased CTCF or cohesin binding. The 3’ portion of *C9* overlaps an antisense lncRNA that shows a much higher male-bias in expression (*lnc12340*; M/F = 26) (Fig. 4C, Fig. S5C). A cluster of strongly male-biased enhancers lies ∼230 kb upstream of the TSS of *C9*, and is characterized by male-biased DHS and H3K27ac peaks, whereas the TSS of *C9* is only comprised of a male-biased DHS. The far upstream enhancer and the TSS of *C9* both fall just outside of (< 10 kb from) a nested intra-TAD loop (Fig. 4C, bottom track) that encompasses *Dab2*, whose expression in male liver is 87-fold lower than *C9* (FPKM = 0.7 vs 61). The promoter region of *C9* interacts with the cluster of far upstream enhancers (Fig. 4C, red arrow), with stronger interactions seen in male liver. Weaker, mostly non-focal interactions were seen between the *C9* promoter and sites within the nested intra-TAD loops. This apparent insulation allows the strong male-biased upstream enhancers to bypass the more proximal *Dab2* and drive expression of *C9*. This insulation-by-looping mechanism also enables the 87-fold higher expression of *C9* compared to *Dab2* in male liver. However, a shorter isoform of *Dab2* contained within the larger nested intra-TAD loop does show weak, male-specific interactions despite a lack of sex-bias in expression. The movement of cohesin along chromatin has been linked, at least in part, to transcriptional activity [19, 61, 62]. Thus, higher levels of transcription in male liver could lead to less pausing of cohesin at loop anchors and thereby increase interactions across these loop boundaries in males. Given the interaction with the far upstream enhancer element, we hypothesized that the expression of *C9* and the antisense *lnc12340* would be sensitive to the loss cohesin of binding. Indeed, both genes showed a 3 to 5-fold decrease in expression upon cohesin depletion in mouse liver (p < 0.01 for both), while the insulated gene *Dab2* showed no significant change in expression (Fig. 4D).

***Nudt7*** is a highly expressed, male-specific gene with distal male-biased enhancers within the same intra-TAD loop (M/F = 2.5; FPKM = 132 in male liver). Although there are some weak male-biased DHS within the gene body of *Nudt7* (Fig. 4E), there is no apparent sex bias at its shared promoter with the sex-biased lncRNA gene *lnc7430* (M/F = 4.0; FPKM = 2.9 in male liver). Approximately 22 kb and 39 kb upstream of the TSS of *Nudt7* are two male-biased enhancers with male-biased DHS and H3K27ac marks; the latter also is proximal (∼2.5 kb upstream) to the TSS of the male-biased *lnc7423* (M/F = 7.0; FPKM = 2.8 in male liver). This enhancer cluster is one of 503 super-enhancers identified in both male and female liver [32]. We observed male-biased 4C-seq interactions between the enhancer viewpoint and a neighboring male-biased enhancer, and also with the shared *Nudt7* and *lnc7430* promoter region (Fig. 4E, red arrows; Fig. S5D). In contrast, we did not observe focal interactions in either sex between the enhancer viewpoint and a sex-independent enhancer 15.7 kb upstream of *Nudt7*. Both *Nudt7* and *lnc7430* were strongly down regulated in male liver upon cohesin depletion (Fig. 4F), suggesting their expression is dependent on interactions facilitated by the CAC-anchored intra-TAD loop that encompasses this genomic region. Expression of *lnc7423* was not significantly reduced, perhaps due to its closer proximity to strong male-biased enhancers.

### Sex-independent, nested intra-TAD loops restrict *Nox4* to proximal enhancer-promoter interactions

NADPH oxidase 4 (*Nox4*) exhibits male-biased expression in mouse liver (M/F = 7.7) and may contribute to a number of liver pathologies whose incidence or severity is male-biased [63, 64]. *Nox4* is highly up regulated in tumor compared to healthy liver tissue of mice that spontaneously develop liver tumors (Fig. S7A), and in humans, *Nox4* is up regulated in hepatocellular carcinoma and other cancers (Fig. S7B). Mouse *Nox4* is located within a pair of nested intra-TAD loops (Fig. 5A, bottom). Mouse *Nox4* also has a strong male-biased enhancer 11.5 kb upstream of the TSS and a strong male-biased DHS 125 kb downstream of the TSS (green vertical highlight). However, only the upstream region has H3K27ac (active enhancer) marks (Fig. 5A). We placed 4C-seq viewpoints at both the upstream region (viewpoint VP1, at −11.5 kb; red highlight in Fig. 5A) and the downstream region (viewpoint VP2, at +125 kb; green highlight) to investigate chromatin interactions with each putative regulatory region. Interactions with the downstream DHS at VP2 were limited to the domain defined by the pair of 3’ anchors of the nested intra-TAD loops, consistent with these loops insulating from distal interactions (Fig. 5A, green bracket at bottom). As a result, VP2 did not interact with the promoter of *Nox4* or with the −11.5 kb enhancer in either male or female liver. Furthermore, VP2 did not show any consistent male-biased interactions, despite its location at a strong male-biased DHS.

**Fig. 5.**
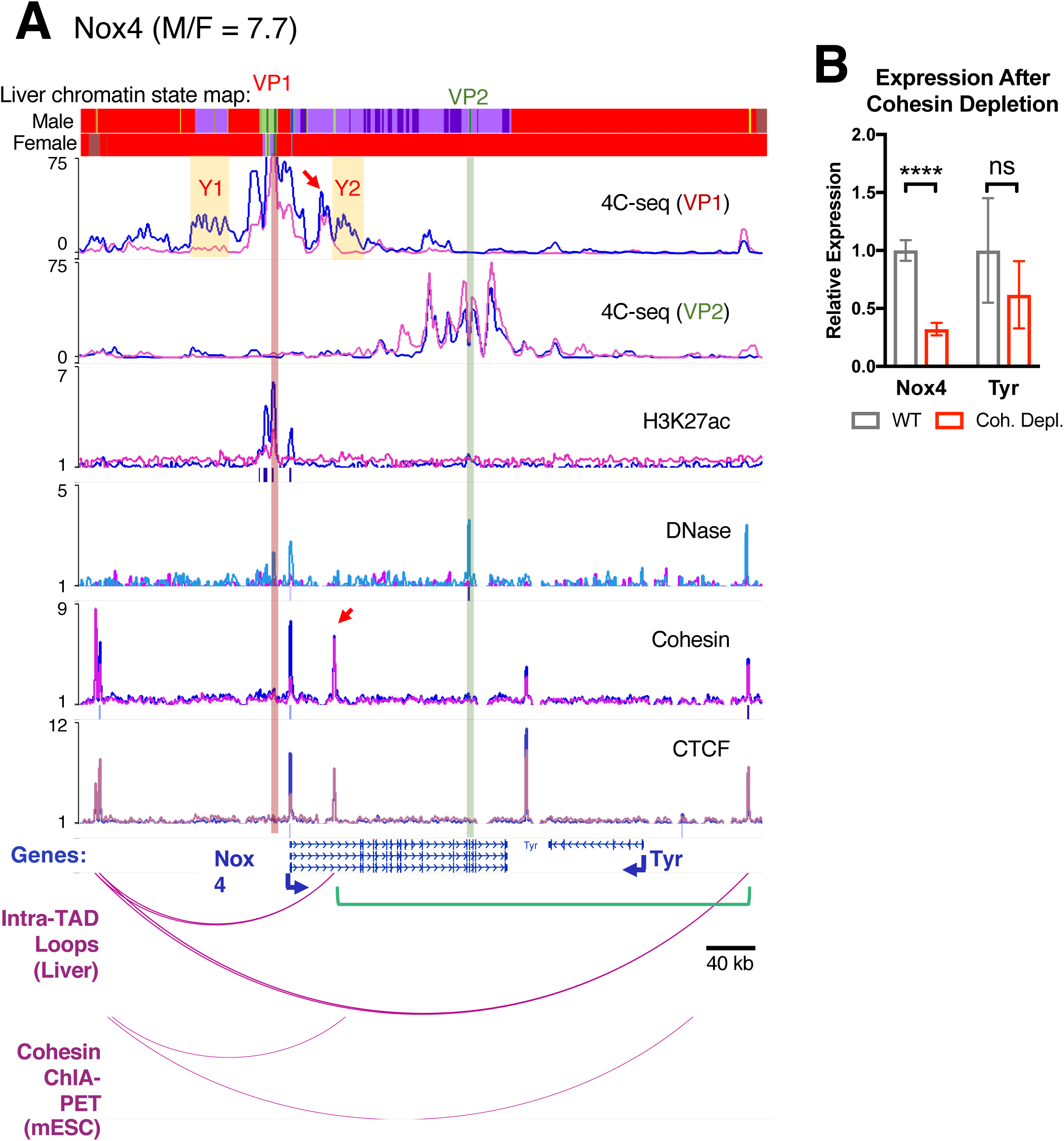
Nox4 results with 4C-seq for two viewpoints and confirming nested intra-TAD loop structure. **A**. Two viewpoints for *Nox4* gene region: VP1 is anchored ∼11 kb upstream of the *Nox4* TSS (light red highlight), and VP2 is anchored ∼125 kb downstream of the TSS (green highlight). VP1 has the strongest male-biased H3K27ac mark. This region shown spans chr7:94248242-94726358. In addition to the ChIP-seq (H3K27ac, Cohesin, and CTCF) and DNase-seq tracks shown, the tracks at the top show the chromatin state of this genomic region in male liver (top) and female liver (bottom). Chromatin states are colored: green indicates an enhancer-like state, blue indicates a promoter-like state, and purple a transcribed-like state. Red indicates an inactive chromatin state (see Fig. S7C for further details). Regions Y1 and Y2 are in chromatin state E13 in male liver but in inactive stateE2 in female liver. Y1 and Y2 both include a short enhancer state region in male liver (E11 within region Y1; E10 in region Y2). The absence of focal interactions between VP1 and VP2 supports the model of two nested and insulated intra-TAD loops shown at the bottom. All tracks were normalized and are presented as described in Fig. 4. **B.** Expression of *Nox4*, but not the neighboring gene *Tyr*, is cohesin-dependent. Although *Nox4* is primarily regulated by proximal enhancers within the shorter intra-TAD loop, its full expression is nevertheless dependent on cohesin. This may be due to the need for the intra-TAD loop structure; however, loss of this insulation did not increase expression of *Tyr*. Expression of *Nox4* was reduced by 62% (p<0.0001). Data presentation is as described in Fig. 4D.

In contrast, the −11.5 kb enhancer at VP1 showed male-biased interactions with several genomic regions, including the *Nox4* promoter and two genomic regions that are in a transcribed-like chromatin state [8] in male liver, but are in an inactive state in female liver (Fig. 5A, regions Y1 and Y2; purple and red genomic segments in chromatin state maps at *top*, respectively). Both VP1-interacting regions include a short segment in an enhancer state (Fig. 5A, green band within purple segment; see Fig. S7C). We also observed 4C-seq interactions between VP1 and a region just upstream of the shorter intra-TAD loop 3’ anchor 31 kb downstream of the *Nox4* TSS in both male and female liver (Fig. 5A, red arrow; Fig. S7D). The high intra-domain interactions that we observed for VP1 and the shorter nested intra-TAD loop, and the low frequency of inter-domain interactions, are indicative of the insulation activity of these loops [32]. Immediately downstream of the shorter nested loop’s 3’ anchor, in region Y2, we observed 4C-seq interactions specific to male liver, which may be due to increased movement of cohesin beyond this loop anchor as a result of the increased transcription of *Nox4* in male liver. Movement of cohesin has been directly linked to transcription both *in vitro* and *in vivo* [19, 61, 62], and may result in a more dynamic loop structure and weaker local insulation at the 3’ loop anchor of the shorter nested loop. The VP1-interacting regions Y1 and Y2 are both in a transcribed chromatin state only in male liver, consistent with the male-biased 4C-seq interactions of VP1 with both regions.

These findings support the predicted model of two nested intra-TAD loops, with the shorter enclosed loop insulated from the larger enclosing loop. Domain predictions for other mouse tissues, based on computational methods and experimentally-observed looping in mouse embryonic stem cells [27, 52], support the conclusion that the genomic regions defined by VP1 and VP2 are in separate domains (Fig. 5A, bottom). Accordingly, only the −11.5 kb enhancer at VP1 would be predicted to interact with the *Nox4* promoter. Generally, the *cis* regulatory elements relevant for the regulation of *Nox4* appear to be contained within the shorter intra-TAD loop. It is less clear what regulatory function the male-biased DHS at VP2 plays, as it does not interact with *Nox4* or with the downstream gene, *Tyr*, which is not expressed in liver (FPKM < 0.01 in both sexes).

## Discussion

We investigated sex differences in autosomal 3D genome organization in the mouse liver model, focusing on sex-based differences in chromatin binding and interactions involving cohesin and CTCF, which mediate long-range DNA looping interactions that segment mammalian genomes into megabase-scale TAD domains and their shorter intra-TAD domains. We identified 1,847 binding sites for cohesin and/or CTCF that show significant differential occupancy between male and female mouse liver; however, very few of these sites were associated with sex differences in TAD or intra-TAD loop anchors. A majority of the sex-biased binding sites classified as cohesin-non-CTCF (CNC) sites (but only a minority of cohesin-and-CTCF (CAC) and Lone CTCF sites) mapped to distal enhancers, and a major subset of these overlapped sex-biased distal enhancers (median distance 238 kb to TSS of a sex-biased gene). These findings are consistent with the general role of cohesin in mediating distal enhancer−promoter interactions [26, 28], and more specifically, indicate a role for sex-biased cohesin binding in sex-biased enhancer activity. We also found that male-biased genes with distal but not proximal sex-biased enhancers were much more sensitive to cohesin depletion than genes with proximal sex-biased enhancers, implicating cohesin in long-range enhancer interactions regulating these sex-biased genes. Finally, by applying 4C-seq to sex-biased enhancer viewpoints in five genomic regions, we established that sex differences in chromatin interactions are a common feature of sex-biased gene expression in the liver, and we elucidated how chromatin interactions link sex-biased genes to distal sex-biased enhancers, guided both directly and indirectly by cohesin and/or CTCF looping.

Although TADs and intra-TADs are largely conserved across tissues, 20-30% of all such CAC-mediated loops are cell type-specific [32]. Nevertheless, when comparing male and female mouse liver, which show extensive growth hormone-regulated differences in epigenetic state [8, 9, 14], we did not find evidence for widespread formation of sex-specific intra-TAD loops. Rather, we found that intra-TAD loops in mouse liver are largely sex-independent and devoid of sex-biased CTCF or cohesin binding at their CAC anchors. These loops do have the ability, however, to indirectly facilitate sex-dependent chromatin interactions. Thus, a sex-independent intra-TAD loop was shown to insulate the super-enhancer-associated male-biased gene *Nudt7*, and nested intra-TAD loops insulating *Nox4* restricted the promoter of this male-biased gene from an intronic enhancer while enabling interactions with a cluster of upstream enhancers. Furthermore, our analysis of female-biased gene regions revealed female-biased proximal enhancer−promoter interactions in the *Sult3a* gene region associated with female-biased cohesin binding, as well as female-biased interactions between the *A1bg* promoter, a far distal (>100 kb) enhancer, and distal female-biased CTCF binding sites. Together, these findings support the proposal that CTCF and cohesin contribute in both direct and indirect ways to the formation of sex-biased enhancer-promoter contacts in mouse liver (Fig. 6).

**Fig. 6.**
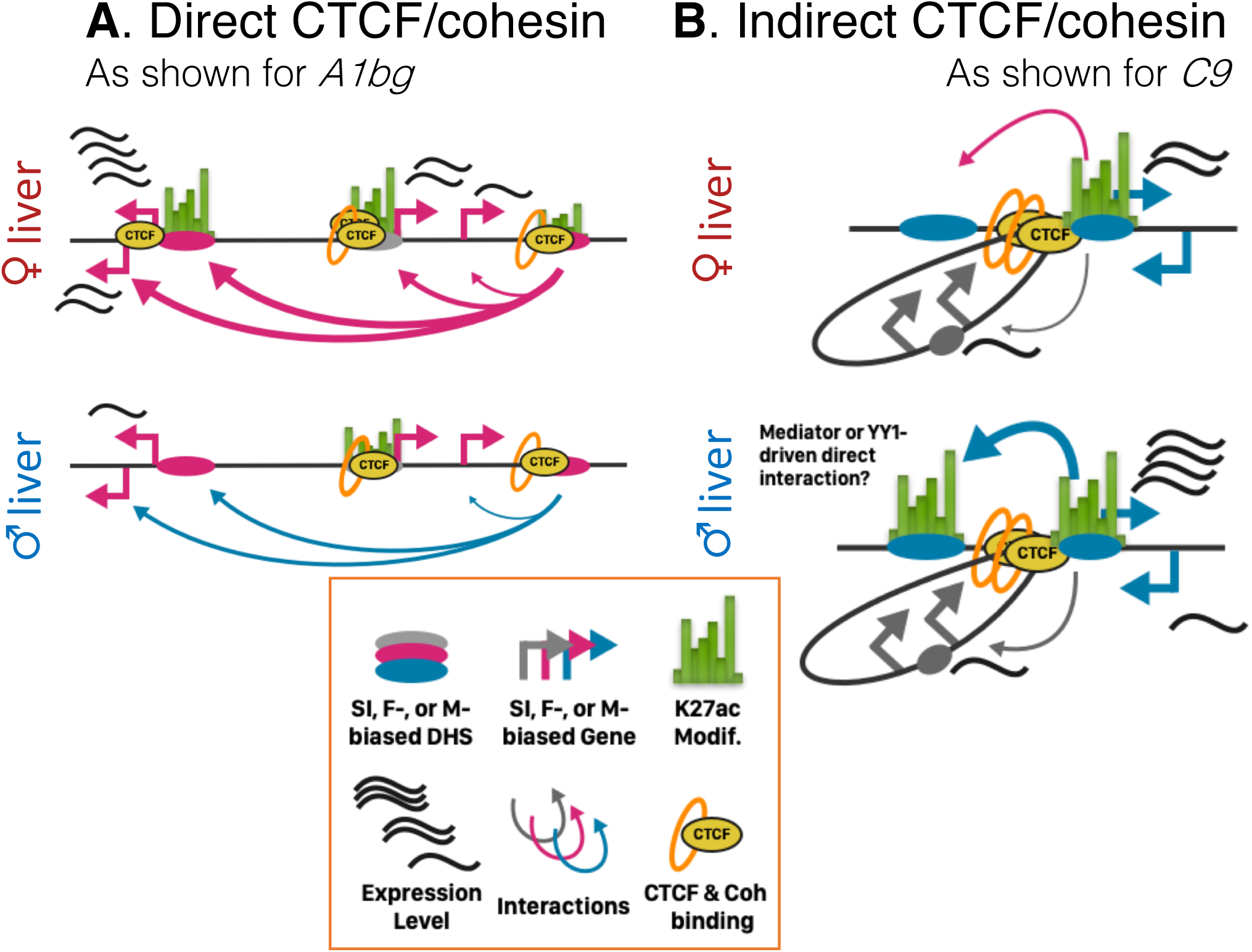
Model of the direct and indirect contribution of cohesin and CTCF to sex-biased liver gene expression. *A1bg* exemplifies direct regulation, where female-biased CTCF binding can explain the observed female-bias in looping interactions. *C9* is an example of intra-TAD loops present in male and female liver that contribute indirectly to sex-biased gene expression. Not only does the intra-TAD loop insulate the lowly-expressed sex-independent gene *Dab2*; it also brings the distal male-biased enhancer into close proximity. Likely additional factors, such as Mediator, YY1, or eRNAs, can contribute directly to the interactions observed in male liver.

Our analysis of the male-biased complement factor gene *C9* provides an interesting example of sex-independent CAC-looping that indirectly facilitates sex-biased enhancer-promoter contacts. *C9* interacts strongly with a distal (>200 kb upstream) male-biased enhancer while bypassing the weakly expressed (and sex-independent) *Dab2* gene region, which is insulated by a nested pair of intra-TAD loops. These nested loops, in turn, bring the TSS of *C9* into much closer proximity of a cluster of far upstream male-biased enhancers than would be achieved based on linear genomic distance alone (Fig. 6B). Furthermore, we observed more frequent contacts between *C9* and the far upstream enhancers in male compared to female liver, despite the absence of any male-biased binding of CTCF or cohesin to help explain sex differences in contact frequency. Conceivably, the male-biased DHS located in the upstream region facilitate direct enhancer−promoter interactions specifically in male liver by working in the context of other known looping factors. Mechanistically, these interactions could be driven by proteins, such as YY1 [55] or Mediator [26], or by non-protein factors such as enhancer-RNAs (eRNAs) [65, 66]. Of note, 3 of the 4 male-biased enhancers far upstream of *C9* are actively transcribed to produce bidirectional eRNA transcripts in male mouse liver [67], and all four regions are bound by the protein YY1 [68] (Fig. S5G).

A majority of the 1,847 sex-biased binding sites for CTCF and cohesin identified here are intergenic and distal to sex-biased genes and enhancers. Furthermore, fewer than half of these sex-biased binding sites are located in TADs with a known sex-biased gene. Although those sites lack any obvious link to sex-biased gene expression in untreated liver, they could have a priming effect and contribute to sex-specific responses reported for hepatic stressors, such as high fat diet [69] and xenobiotic exposure [70], which we recently showed can induce a sex-biased gene response in TADs whose genes do not show a sex-bias in expression in the unstressed state [71]. Just as short-term feeding of a high fat diet can leave a lasting epigenetic memory in the form of epigenetic modifications [72], sex-biased binding of CTCF and/or cohesin may differentially prime each sex for distinct looping patterns that enable the observed sex-biased responses to chemical exposure or dietary stressors.

While CAC sites at TAD and intra-TAD boundaries have a well-established role as anchors that enable loop domain-level nuclear organization [18, 27, 28, 32, 33], non-anchor CAC sites may directly link enhancers to promoters or to other enhancers, and thereby contribute to interactions governing tissue-specific gene expression [36, 42]. A majority of sex-differential CAC binding occurs at non-anchor CAC sites (Table S1), a subset of which may mediate long-distance interactions involving sex-biased enhancers and gene promoters. Specific examples described here include the enhancer−enhancer and enhancer−promoter contacts that we identified by 4C-seq for *A1bg* in the context of female-biased CTCF binding. Similarly, more than half of male-biased liver CTCF binding occurred at Lone CTCF sites, which we found are closer than CAC sites to gene TSS, and can also play a non-canonical role in looping between enhancers and promoters [36].

4C-seq interactions between sex-biased enhancer viewpoints and distal sex-biased lncRNAs [11, 12] were found in three of the five sex-biased genomic regions we investigated. In one example, the highly female-biased gene *A1bg* is nearby three clusters of strongly female-biased, nuclear-enriched mono-exonic lncRNAs, several of which are transcribed from genomic loci that show female-specific interactions with the distal female-specific enhancer viewpoint that we interrogated. Enhancer-associated lncRNAs have been defined as intergenic transcripts with enhancer chromatin marks whose expression is tissue-restricted and is associated with increased expression of nearby expressed protein coding genes [73, 74]. Based on our findings, we propose that functional enhancer-associated lncRNAs might be identified by their looping interactions with enhancer sequences, which can be determined globally using high throughput interaction methods, such as Hi-C [75].

The sex-biased cohesin and CTCF binding sites described here were discovered using livers from adult (8 week) mice, and likely encompass only a subset of all sex-differential cohesin and CTCF binding sites across the lifespan of a mouse, given the dramatic changes in sex-biased gene expression that occur during prenatal and especially postnatal liver development [76, 77]. The sex-biased binding of cohesin and CTCF to liver chromatin in adult liver is expected to be regulated by pituitary growth hormone secretion, which is sex-dependent and produces the sex-dependent plasma growth hormone profiles that regulate the vast majority of sex differences in the adult liver, including differences in gene expression [14, 78], transcription factor binding [79, 80], and chromatin states [8, 9, 14]. Given that CTCF binding to DNA can be inhibited by DNA methylation [48, 49, 81], the hypomethylation of enhancer sequences seen in male compared to female liver [50] could contribute to male-specific CTCF binding at such sites. Such an effect is expected to become more pronounced with age, given the increased male hypomethylation reported in older mice [50, 82].

The sex-specific patterns of pituitary growth hormone secretion regulating sex-differences in the liver emerge at puberty, and have been implicated in the dynamic regulation of liver chromatin states in both male and female adult mouse liver [9, 14]. We do not know when during mouse liver development the sex-differential chromatin interactions described here are first established, whether they constitute a relatively fixed (static) 3D framework governing transcription in male and female nuclei, or alternatively, whether they respond dynamically to the temporal changes in plasma growth hormone profiles that regulate sex differences in liver chromatin states. The potential for dynamic, reversible changes in DNA looping was demonstrated in a study where auxin-induced cohesin degradation led to a loss of virtually all DNA loops after 40 min, followed by their reestablishment within one hour of auxin withdrawal and cohesin reintroduction [23]. Extensive changes in chromatin interactions also occur during circadian oscillations in mouse liver [83-85] and during macrophage differentiation [86]. Our finding that the expression of distally-regulated male-biased genes is highly sensitive to cohesin depletion illustrates the importance of distal enhancer−promoter interactions in maintaining male-biased gene expression. This may not be a direct result of a loss of sex differential cohesin and/or CTCF binding at distal male-biased enhancers, as only 5 of 32 distally-regulated male biased genes show male-biased CTCF/cohesin binding at distal male-biased enhancers (data not shown). While the sex-biased enhancer−promoter and enhancer−enhancer loops that we describe here might be dynamically regulated by growth hormone, they could also be determined by intrinsic sex differences [78], or might be established by early postnatal hormone exposures that program liver gene expression [87]. Studies to detect dynamic changes in DNA looping and quantify changes in chromatin interaction strength, e.g., during the course of a male plasma growth hormone pulse [14], will likely require technical improvements to the 4C-seq protocol. These include the use of unique molecular identifiers for more accurate quantification of interactions [60] and the elimination of any PCR artifacts associated with over-amplification, which are difficult to address using conventional 4C-seq methods [47] and may have decreased the magnitude of the apparent sex differences in chromatin interactions seen in our work.

## Conclusions

We employed the mouse liver model with its extensive sex differences in gene expression to study sex differences in nuclear organization and DNA looping interactions in a non-reproductive tissue exposed to sex-unique patterns of hormone stimulation. We determined that male-biased genes with distal but not proximal sex-biased enhancers are particularly sensitive to the loss of cohesin binding. Furthermore, while most sex-biased binding sites for CTCF and cohesin were found to be distal from sex-biased genes, a subset likely contributes to sex-biased looping between regulatory elements in *cis*, as exemplified by the female-biased DNA looping interactions observed for *A1bg*. In addition, sex-independent CAC-looping may indirectly provide sex specificity to chromatin interactions by insulating male-biased genes such as *Nudt7*, or by bringing a sex-biased gene into closer proximity to a cluster of sex-biased enhancers, as demonstrated for *C9*. Together, these findings illustrate the direct and indirect contributions that cohesin and CTCF can make to sex-biased gene expression in the liver, and may be broadly applicable to other biological systems where distal regulation of gene expression is of interest.

## Methods

### Mouse protocols, extraction of liver nuclei, and chromatin preparation

Male and female CD-1 mice were purchased from Charles River Laboratories (strain # Crl:CD1(ICR)) and housed separately in the Boston University Laboratory Animal Care Facility. All animal protocols were specifically reviewed for ethics and approved by the Boston University Institutional Animal Care and Use Committee. Mice were euthanized by cervical dislocation at 8 weeks of age, livers were excised, and nuclei were purified then crosslinked with 0.8% formaldehyde for 9 min at 30°C [32]. The crosslinked chromatin was sonicated and stored at −80°C after a small aliquot was removed for crosslink reversal (6 h at 65°C in a thermocycler with a heated lid) to quantify the chromatin by Quanti-iT PicoGreen assay (Invitrogen, cat. # Q33130) and determine the fragment size distribution, as described [32].

### Chromatin immunoprecipitation and sequencing (ChIP-seq)

Immunoprecipitation of sonicated mouse liver chromatin was performed as described [8]. Specifically, 5 µl of rabbit polyclonal antibody to either CTCF (Millipore, cat. # 07-729) or to the cohesin subunit Rad21 (Abcam, cat. # ab992) was mixed with 30 µl of Protein A Dynabeads (Invitrogen, cat. # 1002D) and incubated in blocking solution (0.5% bovine serum albumin in PBS) for 3 h at 4°C. Beads were then washed with blocking solution, followed by incubation overnight with 70 µg of sonicated liver chromatin in 1X RIPA buffer (50 mM Tris-HCl, pH 8.0, 150 mM NaCl, 1% IPEGAL, 0.5% deoxycholic acid) containing 0.1% SDS. After washing with 1X RIPA (containing 0.1% SDS), formaldehyde crosslinks were reversed for 6 h at 65°C, followed by RNase A digestion (Novagen, cat. # 70856) at 37°C for 30 min and then proteinase K digestion (Bioline, cat. #37084) for 2 h at 56 °C. The resulting crude DNA extract was purified using a QIAquick Gel Extraction Kit (Qiagen, cat. # 28706) and quantified with a Qubit HS DNA kit (Invitrogen, cat. # Q32854). All samples were processed using the same protocol and conditions. Sequencing was performed for a total of eight CTCF ChIP-seq samples (n=4 individual male and n=4 individual female livers) and a total of six Rad21 ChIP-seq samples (n=3 male, n=3 female livers). The male liver ChIP-seq samples were those reported previously [32] and are available at GSE102997.

### Impact of cohesin depletion sex-biased gene expression

RNA-seq data for wild-type and cohesin depleted (Nipbl-deficient) mouse liver (GSE93431) [22]. was analyzed for n=4 wild-type (control) and n=4 Nipbl-depleted male mouse liver replicates. Data was FPKM-normalized, and reads were expressed relative to the mean of the wild-type group, which was set = 1 on a per gene basis by dividing the expression value for each individual replicate by the mean of the wild-type group plus a small pseudo count (1e-6) to avoid dividing by zero. Data is presented as mean relative expression ± SD for all plots. To determine the global effects of cohesin depletion on male-biased genes, the set of all expressed, strongly male-biased genes (FPKM > 1, and male/female (M/F) expression ratio > 3) was divided into two groups based on distance from their TSS to the nearest male-biased DHS or male-biased H3K27ac-marked region. Male-biased genes with proximal sex-biased enhancers (n=29) were defined as having their TSS < 20 kb from the nearest male-biased DHS *and* from the nearest male-biased H3K27ac peak; and male-biased genes with distal sex-biased enhancers (n=32) were those with TSS > 20 kb from both such regions. The underlying expression values for all genes are provided in Table S2A and Table S2B.

### 4C-seq methods

Isolation of liver nuclei and crosslinking were performed as described [32] through the step where crosslinked nuclei were pelleted by centrifugation. Digestion, proximity ligation, and inverse PCR were then carried out as described [32]. Briefly, frozen nuclei were resuspended in Buffer A (15 mM Tris-HCl pH 8.0, 15 mM NaCl, 60 mM KCl, 1 mM EDTA pH 8.0, 0.5 mM EGTA pH 8.0, 0.5 mM spermidine, 0.3 mM spermine) and quantified using a Countess Automated Cell Counter (Invitrogen, cat. # C10227). An aliquot of 10 million nuclei was used for each individual 4C experiment. Nuclei were pelleted at 1,000 x g for 5 min at 4°C then resuspended in 450 μl of 1X NEBuffer 3 (NEB, cat. # B7003S). Primary restriction enzyme digestion of intact nuclei was carried out overnight at 37°C using 50,000 U of DpnII (NEB: #R0543) with agitation at 900 RPM. DpnII was inactivated by adding SDS to a final concentration of 2%. Samples were then diluted 5-fold in 1X ligation buffer (Enzymatics, cat. # B6030). Proximity ligation was performed overnight at 16°C with 200 U of T4 DNA ligase (Enzymatics, cat. # L6030). DNA was then reverse-crosslinked and purified using a standard phenol/chloroform cleanup method following the manufacturer’s protocol (VWR, cat. # VWRV0883). Secondary digestion of purified DNA was performed overnight at 37°C with 50 U of Csp6I (Thermo Scientific, cat. # ER0211) in 500 µl of 1X Buffer B (Fermentas, cat. # BB5; 10 mM Tris-HCl (pH 7.5), 10 mM MgCl_2_, 0.1 mg/ml BSA), followed by heat inactivation at 65°C for 30 min. Samples were diluted 10-fold and secondary ligation was carried out overnight at 16°C, as described above. The effectiveness of primary digestion, proximity ligation, secondary digestion, and secondary ligation was verified at each step by reverse crosslinking and gel electrophoresis analysis (see gel image in Fig. S6A). The final PCR template was purified by phenol/chloroform clean up, followed by QiaPrep 2.0 column cleanup (Qiagen, cat. # 27115) to yield a standard, circularized 4C inverse PCR template, which was amplified using specific viewpoint primers, as described below.

PCR reactions were performed using inversely oriented primer pair sequences for valid 4C-seq viewpoints, obtained from the 4CSeqpipe primer database [http://www.wisdom.weizmann.ac.il/~atanay/4cseq_pipe/], with the addition of 5’ dangling truncated Illumina adapters (Table S3A). Candidate viewpoints were selected based on the following criteria. First, we only considered viewpoints that are in the same TAD as at least one protein-coding or lncRNA gene showing >3-fold sex bias in its expression. Second, the viewpoint must be within 1 kb of the transcription start site (TSS) of a sex-biased gene, or it must overlap a sex-biased enhancer (minimum 2-fold sex-bias in the sex bias in normalized DHS opening or H3K27ac mark intensity). Third, the non-reading primer (Table S3A) was required to map to the genome uniquely, while the reading primer was more stringently required to have > 89% unique sequence identity (i.e., no 18-mer within a 20 nt primer sequence that maps elsewhere in the genome). Inverse PCR amplification of 1 microgram of each 4C template was performed using Platinum Taq DNA polymerase (Invitrogen, cat. #10966026), as follows: 94°C for 2 min, 25 amplification cycles (94°C for 30 s, 55°C for 30 s, 72°C for 3 min), then 4°C hold. To minimize the impact of PCR artifacts, a total of 6 identical inverse PCR reactions were performed for each sample, except as noted. These replicate reactions were processed in parallel and pooled prior to library preparation. For 4C-seq analysis of the *Nox4* genomic region viewpoints, single independent inverse PCR reactions were carried out for each of four liver samples (males M1 and M2; females F1, and F2) and sequence libraries were then prepared independently. Sequencing data was pooled at the raw read level. Examples of pooled 4C libraries are shown in Fig. S6B.

4C-seq samples were amplified using barcoded primers, such that each biological replicate had a unique barcode (NEB, cat. # E7335). To minimize over-amplification, each sample was amplified with 5 additional cycles of PCR to fill in the full Illumina adapter sequence needed for sequencing. Due to the low sequence complexity in the first 20 bases of each sequence read, 4C-seq libraries were multiplexed by combining with high complexity sequencing libraries for unrelated samples (e.g., RNA-seq or ChIP-seq libraries), which were sequenced in the same Illumina sequencing lane. In practice, 4C-seq libraries constituted no more than 15% of the total library pool, by molarity. Samples were sequenced on an Illumina HiSeq 2500 instrument for 50, 125, or 150 bp paired end reads.

### Computational analysis of ChIP-seq datasets

Sequence reads were split by barcode and mapped to mouse genome assembly mm9 using Bowtie2 (v2.2.9). All reads not uniquely mapped to the genome were excluded from downstream analyses. Peak calling was performed using MACS2 (v2.1.1) with default parameters, and peaks that overlapped blacklisted genomic regions (www.sites.google.com/site/anshulkundaje/projects/blacklists) were filtered out. Additionally, we removed spurious peaks that exclusively contained PCR duplicated reads, defined as 5 or more identical sequence reads that do not overlap any other reads. All BigWig tracks used to visualize sequencing data in a genome browser were normalized for both sequencing depth and sample quality, expressed as reads in peaks per million mapped reads (RIPM). In practice, the browser y-axis displays the read count from a given sample divided by the total number of reads in peaks (reads that overlap a peak identified in any sample, or the union peak list), per million. Normalization was performed separately for the CTCF and cohesin datasets. This approach is functionally similar to the quality control metric known as Fraction of Reads in Peaks (FRiP) used by the ENCODE consortium [88]. These analyses were facilitated by a ChIP-seq analysis pipeline described elsewhere [89]. All samples used in this study were judged to be of good quality, with a mean FRiP value of 0.217 and ranging from 0.103 to 0.344. A full listing of samples sequenced and sequencing statistics is provided in Table S3B.

To identify sex-differential ChIP-seq peaks, diffReps (v1.55.4) [90] was used with default parameters and a window size of 200 bp to identify in-peak differential sites, i.e., diffReps sites that overlap a MACS2 peak, defined below. The diffReps output list of sites was filtered to remove diffReps-identified sites that did not meet the following conditions: overlap with at least one of the peaks in the union peak list for the relevant factor (Table S1F, Table S1G for CTCF; Table S1H, Table S1I for cohesin), contains at least 10 sequence reads, shows >2-fold sex difference, and has an FDR < 0.05. The resultant sets of ChIP-seq peaks were defined as standard stringency sex-biased peaks, and were used for all analysis, except as noted. A set of lenient stringency sex-biased CTCF and cohesin peaks was defined, as follows. Sequence reads for biological ChIP-seq replicates were combined (merged) to give a single merged sample for each sex and each factor (Male CTCF, Male cohesin, Female CTCF, and Female cohesin). For each transcription factor, the male and female merged samples were then compared, and the merged sample with the higher FRiP was down-sampled, so that the mapped read files for each sex contained the same, normalized number of reads in peaks. In practice, the combined mapped reads for the merged female cohesin samples, and for the merged male CTCF samples, were down-sampled by a factor of 0.979869 and 0.745955, respectively. MAnorm [91] was then used to compare the FRiP-normalized male and female samples to identify a set of sex-differential binding sites for each factor (minimum 2-fold sex bias; p-adj < 0.01; read count > 15 for the up-regulated peak). Binding sites identified by MAnorm that were not on the standard sex-biased peak list were designated lenient stringency sex-biased peak lists. Bedtools (v2.26.0) was used for overlap analysis to determine the distance of ChIP-seq peaks to other genomic features. Genomic coordinates (mm9) for TAD and intra-TAD boundaries were downloaded from [32], where TAD definitions are based on experimental Hi-C analysis for male mouse liver [35].

Sex-differential ChIP-seq peaks were analyzed separately for the sets of male and female biological replicates and then divided into four groups, CAC(ΔCoh), CAC(ΔCTCF), CNC(ΔCoh), and Lone CTCF(ΔCTCF), based on the following criteria. Sex-differential cohesin peaks (ΔCoh) were designated CAC(ΔCoh) peaks if they overlapped a CTCF peak identified in at least 3 of the 4 CTCF ChIP-seq biological replicates in the sex showing higher cohesin binding (e.g., male-biased cohesin peaks were compared to male liver CTCF ChIP-seq replicates). Alternatively, they were designated CNC(ΔCoh) peaks if they overlapped a CTCF peak found in either 0 or 1 of the four CTCF-seq biological replicates. Those ΔCoh peaks that overlapped 2 of the 4 CTCF replicates were excluded from downstream analyses. Similarly, sex-differential CTCF peaks (ΔCTCF) were designated CAC(ΔCTCF) peaks if they overlapped a cohesin peak found in 2 or 3 of the three available cohesin ChIP-seq biological replicates. Sex-differential CTCF peaks were designated Lone CTCF(ΔCTCF) peaks if they overlapped peaks in either 0 or 1 of the three cohesin ChIP-seq biological replicates. 137 sex-differential CTCF peaks overlapped sex-differential cohesin peaks, and were thus CAC(ΔCoh/ΔCTCF); 50 of these 137 CAC peaks were autosomal (Table S1C). Sex-independent cohesin peaks, and sex-independent CTCF peaks, were respectively defined by ranking each peak based on the following ratio: (RIPM-normalized ChIP signal for the merged male sample) / (RIPM-normalized ChIP signal for the merged female sample), performed separately for CTCF and for cohesin. The 1,000 peaks whose ratios were closest to 1 were defined as the set of sex-independent CTCF, and cohesin, peaks.

### Discovery of intra-TAD loops

CTCF motif discovery was performed using the FIMO option from MEME Suite (v4.10.0), and presence of a motif was defined as a motif score > 10. Intra-TAD loops for female mouse liver were identified using the computational method described previously for male liver [32]. The analysis pipeline was run for all CAC sites and with an initial loop count of 20,000, using the set of default parameters reported for male liver [32]. This analysis yielded 9,724 intra-TAD loops with 10,273 loop anchors in female liver; this compares to 9,543 intra-TAD loops and 9,052 loop anchors identified in male liver [32]. The redundancy in loop anchors is a reflection of nested CAC-mediated loop structures, as was described in other studies using experimentally-measured loop identification compared to the computational approach used here; these studies include ChIA-PET analysis of the cohesin subunit SMC1A in mouse embryonic stem cells [27] and Hi-C analysis in human GM12878 cells, where 9,448 loops were associated with 12,903 loop anchors [18, 33]. Reciprocal overlap between loops was analyzed using bedtools (bedtools intersect –wa –u –r –f 0.8), as described [32]. A total of 2,527 intra-TAD loops were unique to either male or female liver; however, very few had anchors that overlapped a sex-differential CAC site, suggesting that most are not biologically relevant. Supporting this, the loops that were unique to either male or female liver were weaker than the loops shared between male and female livers, and in many cases the loops narrowly met the significance cutoff in one sex but not the other. This finding is similar to our earlier finding that tissue-specific loops are often weaker than those predicted in multiple tissue types [32]. Intra-TAD loops for male and female mouse liver are listed in Table S1J, and female intra-TAD loop anchors are listed in Table S1K; a comparable listing for male liver is available in [32].

### Computational analysis of 4C-seq datasets

Biological replicates were demultiplexed by index read barcode. As the fastq files for each biological replicate contained sequence reads from multiple viewpoints, the reads in each file were further split based on matches to the reading primer for each viewpoint (Table S3A). Then, prior to mapping, we used FASTX-Toolkit (v0.0.14) to remove the first 20 nt of sequence from the 5’ end of each read, as this represents the reading primer. For read length consistency, we trimmed 25 nt from the 3’ end of 150 nt read libraries, making them identical in length to the 125 nt libraries (i.e., both were 105 bases long after 5’ and 3’ trimming). Bowtie2 (v2.2.2) was then used to iteratively map the reads to a reduced mouse genome, which was comprised of all genomic sequences 105 bp upstream and 105 bp downstream from each DpnII cut site in the genome (recognition sequence: GATC), as in [47, 92]). To implement this step, we scanned the genome for all occurrences of the DpnII cut site GATC using the UCSCutils tool oligoMatch (--exactmatch; default parameters). The resulting set of coordinates was then expanded to include sequence 105 bp upstream and 105 bp downstream of each DpnII site (bedtools flank -l 105 -r 105), which was extracted from the mm9 genome using bedtools getfasta. This reduced mouse genome sequence was indexed using bowtie2 prior to mapping. An iterative mapping strategy was implemented because some reads contained multiple ligation junctions, rendering the full-length read unmappable, as described in some Hi-C pipelines [93]. First, non-uniquely mapped reads were trimmed by ∼10% of their total length (starting from the 3’ end of the read), and mapping was reattempted as above. This process was repeated until a minimum read length of 20 nt was reached, with a step size of 2 nt (for 50 nt-long reads) or a step size of 10 bp (for 125 or 150 nt long reads) per iteration. These iterative trimming and remapping steps increased the overall percentage of uniquely mapping reads by 2-6% for shorter read lengths libraries, and by 8-22% for longer read length libraries.

After mapping, bedGraph files were smoothed using ucscutils (v. 20130327) with a sliding window of 11 restriction fragments, taking the median value in the window. Smoothed BigWig tracks were normalized by total reads per million to account for differences in sequencing depth. Merged replicates shown in the main figure panels were generated by taking the median signal at a given restriction fragment per viewpoint and sex. The genomic region overlapping the viewpoint fragment was removed prior to BigWig generation for merged replicates. Tracks were visualized in the WashU genome browser with replicates overlaid using the Matplot feature. Table S3C shows the sequences used for demultiplexing 4C libraries as well as basic statistics for read mapping and quality control.

### Analysis of other datasets

Pan-cancer human expression data for *Nox4* was obtained for tissues with matched Normal and Tumor expression values and analyzed using the web interface TIMER using default parameters (https://cistrome.shinyapps.io/timer/) [94]. *Nox4* expression for male C3H mouse liver and neoplasms was obtained from [95] as processed, normalized RNA-seq counts (Accession # E-MTAB-6972). The following male and female mouse liver datasets were used to generate browser screenshots: DNase-seq [15], male liver CTCF and cohesin ChIP-seq data [32], sex-biased lncRNAs [11], M/F expression ratios [9], and H3K27ac ChIP-seq data [96]. Unless otherwise noted, FPKM and M/F expression ratios for protein coding genes are based on ribosomal RNA-depleted total liver RNA, while lncRNA expression ratios are based on ribosomal RNA-depleted liver nuclear RNA. Bigwig files were RIPM-normalized separately for each experiment type as described above. Genomic regions with significant sex bias are marked by short horizontal bars below each DHS and H3K27ac BigWig signal track. The bars were colored blue to indicate male-biased DHS or H3K27ac regions, and pink to indicate female-biased DHS or H3K27ac regions. A darker shade of color was used to indicate the stringency for the feature: DHS were shaded from dark to light to indicate high, standard, and low stringency for male-biased (blue) and female-biased (pink) DHS, as defined previously [15]. The high and low stringency designations used in that earlier study approximate the stringent and lenient definitions used here. For the sex-biased H3K27ac tracks, a total of four non-overlapping groups were defined based on the magnitude of the fold-change difference in H3K27ac peaks between male and female liver samples (M/F or F/M > 2.5 was defined as strict; and M/F or F/M > 1.5 but < 2.5 were defined as lenient). Both strict and lenient sex-biased H3K27ac peaks were defined using MAnorm with cutoffs of p-adj < 0.05 and read count > 15 for the up regulated peak [91]. These cutoffs resulted in 1,583 female-biased H3K27ac peaks (380 strict, 1203 lenient) and 2,241 male-biased H3K27ac peaks (604 strict, 1637 lenient).

### Statistical analysis

Boxplots, cumulative distribution plots, and statistical analyses were implemented using GraphPad Prism 7. All boxplots are displayed using the Tukey convention, where interquartile range (IQR) was calculated as the difference between the 25^th^ and 75^th^ percentile. Outliers were considered as values falling above the 75^th^ percentile value + 1.5*IQR, or below the 25^th^ percentile value – 1.5*IQR. Whiskers indicate the maximum and minimum values in a set (if no outliers) or from the 75^th^ percentile value + 1.5*IQR to the 25^th^ percentile value – 1.5*IQR (if outliers exist). Boxes indicate the IQR with a horizontal line indicating the median value. Bar graphs show the mean and standard deviation. Unless otherwise indicated, all pairwise comparisons used two-tailed, nonparametric t tests. Comparisons of distributions were performed using a Kolmogorov–Smirnov (K-S) test, while comparisons of values used a Mann-Whitney (M-W) test. The results are annotated in individual figures as follows: **** indicates p ≤ 0.0001; *** p ≤ 0.001; ** p ≤ 0.01; * p ≤ 0.05; and not significant (ns) for p > 0.05.

### Data availability

CTCF and cohesin (Rad21) ChIP-seq data are available under accession numbers GSE130908 for female liver (https://www.ncbi.nlm.nih.gov/geo/query/acc.cgi?acc=GSE130908) and GSE102997 for male liver (https://www.ncbi.nlm.nih.gov/geo/query/acc.cgi?acc=GSE102997). 4C-seq data are available under accession number GSE130911 (https://www.ncbi.nlm.nih.gov/geo/query/acc.cgi?acc=GSE130911).

## Abbreviations

3D: 3-dimensional
4C-seq: circularized chromosome conformation capture-based sequencing
ΔCoh, ΔCTCF: sex-differential binding site for cohesin or for CTCF, respectively
CAC: cohesin-and-CTCF site
CNC: cohesin-non-CTCF site
CTCF: CCCTC-binding factor
DHS: DNase hypersensitive site
FDR: false discovery rate
FPKM: fragment per kilobase per million sequence reads
FRiP: Fraction of Reads in Peaks
K-S: Kolmogorov–Smirnov test
lncRNA: long non-coding RNA
Lone CTCF site: site bound by CTCF but not by cohesin
M-W: Mann-Whitney test
M/F: male/female liver expression ratio
TAD: topologically associating domain
TSS: transcription start site
VP: 4C-seq viewpoint

## Declarations

### Ethics approval and consent to participate

All animal protocols were specifically reviewed for ethics and approved by the Boston University Institutional Animal Care and Use Committee (protocol 16-003).

### Consent for publication

Not applicable

### Availability of data and material

The datasets generated and/or analyzed during the current study are included in this published article and its supplementary information files. Raw and processed sequencing files generated in this study are available at GEO (https://www.ncbi.nlm.nih.gov/geo/) under accession numbers GSE130908, GSE102997 and GSE130911. Other GEO datasets used in our analyses are cited in the text.

### Authors’ contributions

BJM and DJW conceived and designed the study. BJM carried out all of the laboratory experiments and data analyses and prepared the figures and tables for publication. BJM and DJW jointly wrote the manuscript and DJW edited the final manuscript for publication.

## Acknowledgements

We thank Dr. Dana Lau-Corona for sharing enhancer chromatin mark datasets and Dr. Andy Rampersaud for developing the ChIP-seq analysis pipeline used in this study.

## Authors’ information (optional)

None

## Supplemental figures

**Fig. S1 − Additional features of sex-biased differential CTCF/cohesin peaks**

**A.** The limited overlap of sex-biased CTCF and sex-biased cohesin binding, seen in Fig. 1B, is not an artifact of the thresholds used for peak filtering. In the data shown here, similar to Fig. 1B, the overlap of sex-biased CTCF and sex-biased cohesin binding sites is limited. Shown are Venn diagrams with the number of overlapping sex-biased cohesin (*Left*; blue) and CTCF sites (*Right*; purple) for the following groups (top to bottom): (1) All sex-differential sites output by diffReps without filtering for overlap of a MACS2 ChIP-seq peak; (2) All sex-differential diffReps sites (rather than *peaks,* which may contain multiple sites) from diffReps that overlap a MACS2 peak for a given factor; (3) All sex-differential hotspots identified by diffReps, which is an alternate method in the diffReps software package to identify differential sites. Specifically, this last approach looks for clusters of differentially co-regulated sites that might be missed by simple overlap analysis. Overlap for all Venn diagrams is defined as ≥ 1 bp overlapping using bedtools. In some cases, two Rad21 peaks overlap a CTCF peak, or vice versa, and therefore, the number of overlapping cohesin (Rad21) sites does not necessarily equal the number of CTCF sites (hence, two numbers in the Venn overlap).

**B.** Pie charts showing the class distribution of each sex-biased CTCF/cohesin peak set (from top to bottom): male-biased ΔCohesin peaks, female-biased ΔCohesin peaks, male-biased ΔCTCF peaks, and female-biased ΔCTCF peaks. For each of these four groups, the fraction of peaks at CAC sites is shown in purple while the fraction of peaks at either CNC (for ΔCohesin) or Lone CTCF (for ΔCTCF) is shown in blue. The total number of differential peaks in each group is indicated below each chart. Overall, female-biased sites are comprised of a higher percentage of CAC sites than male-biased sites. Consequently, a larger percentage of male-biased peaks are CNC peaks (for ΔCohesin peaks) and Lone CTCF peak (for ΔCTCF peaks). Peak numbers here differ slightly from Fig. 1B for cohesin differential peaks, but not CTCF, because of our approach to categorizing peaks as CNC or CAC for cohesin peaks (see Methods). For CTCF we defined CAC peaks as genomic regions bound by CTCF that were also bound by cohesin in a majority of individual cohesin replicates (2 or 3 out of a total n=3 per sex). Using the same approach for cohesin, we defined CAC peaks as genomic regions bound by cohesin that were also bound by CTCF in a majority of individual CTCF replicates (3 or 4 out of a total n=4 per sex). If a peak was bound by none or only a minority of replicates for the opposite factor then it was considered Lone CTCF (in the case of CTCF; 0 or 1 cohesin replicates overlapping) or CNC (in the case of cohesin; 0 or 1 CTCF replicates overlapping). As CTCF has n=4 replicates, if a cohesin peak is bound by exactly 2 individual CTCF replicates (of the same sex) then it is not classified and is excluded from downstream analyses. 54 male-biased cohesin peaks overlap 2 male CTCF replicates and 36 female-biased cohesin peaks overlap 2 female CTCF replicates (value of 2 in column H of Table S1B). All overlaps were performed using bedtools with a minimum overlap of 1 bp, and all comparisons were made separately for males and females.

**Fig. S2 − Comparison of sex-biased CTCF/cohesin peaks**

**A.** Female-biased CTCF and cohesin peaks tend to be stronger than male-biased peaks. Shown here are box plots for ChIP-seq signal for CTCF and cohesin for both ΔCohesin and ΔCTCF peaks. These plots differ from those presented in Fig. 2A, which present normalized ChIP signal for the factor with differential signal (i.e. male and female cohesin ChIP-seq signal for ΔCohesin peaks). In aggregate, CAC peaks with significant sex-biased cohesin binding show the same directionality of sex-bias for CTCF (and vice versa), albeit at a reduced magnitude (see also Fig. 1C). The y-axis shows normalized ChIP-seq signal for the groups indicated along the x-axis. Peaks with male-biased and female-biased cohesin binding (*Left*) and CTCF binding (*Right*) are presented separately. Each group of 4 box plots represents the male and female ChIP-seq signal for cohesin, followed by the corresponding ChIP-seq signals for CTCF for the same set of peaks. Each plot represents all differential peaks for a given sex (male or female) and factor (CTCF or cohesin). These four datasets are further divided by peak type (CAC or CNC for ΔCohesin peaks, and CAC or Lone CTCF for ΔCTCF peaks), as indicated below the x-axis. Peak scores are calculated by average intra-peak ChIP signal, normalized by the total sequence reads per million in peak (RIPM; see Methods).

**B.** Female-biased CAC peaks contain higher quality CTCF motifs than male-biased CAC peaks [p=0.0212 for CAC(ΔCoh) and p=0.0023 for CAC(ΔCTCF) peaks; M-W t-test], as reflected by the FIMO motif score. This log-likelihood ratio score is a reflection of how close the best intra-peak motif matches the canonical core CTCF motif MA0139.1. There is no significant difference between motif scores for male-biased and female-biased Lone CTCF, or for male-biased and female-biased CNC peaks (p=0.7671 and p=0.1329; M-W t-test). The dashed line at FIMO score = 10 reflects the cutoff used to define the presence or absence of a motif in Fig. S2C.

**C.** CTCF Motif frequency, based on presence of CTCF motif (MA0139.1) as identified by FIMO, with a minimum score of 10. The y-axis shows the percent of peaks in each group (separately for male-biased, female-biased, and sex-independent) found to have a CTCF motif within the peak region. A larger fraction of female-biased than male-biased CAC peaks was found to contain a CTCF binding motif. In contrast, a larger fraction of male-biased Lone CTCF peaks contain a CTCF motif, despite no significant difference in peak strength between male-biased and female-biased Lone CTCF peaks. A larger fraction of female-biased CNC peaks contain a CTCF motif, however, the vast majority do not contain CTCF motifs, as expected (< 20% for all groups). In all cases, the percent for each group is comparable to a matched set of sex-independent peaks.

**D.** Female-biased intra-TAD on Chr19 that contains 12 sex-biased genes, shown in blue and red boxes, some of which are lncRNA genes (ncRNA gene designations, in green). Inset at bottom left of the figure shows CTCF and cohesin (Rad21) ChIP-seq tracks for male and female mouse liver surrounding the 3’ boundary of the female-biased intra-TAD.

**E.** Intra-TAD loops and loop anchors are mostly shared between male and female mouse liver. Using a computational intra-TAD loop prediction algorithm [32], we used the cohesin and CTCF ChIP-seq datasets for male and female mouse liver to identify 9,543 intra-TAD loops in male liver [32] and 9,724 loops in female liver, respectively. 87.9% of the intra-TAD loops in male liver were also identified in female liver (*left)*, and 93.4% of the male intra-TAD loop anchors are also predicted to be loop anchors in female liver. This finding is consistent with there being a limited number of autosomal CAC peaks with sex differences in CTCF and cohesin binding (53 total) (Table S1C) To account for nested loop structures, shared loops were defined as loops with a reciprocal overlap of 80% or greater between the loops, as implemented in prior studies of CAC-mediated insulating loops [27, 32].

**Fig. S3 − Tissue conservation of liver sex-differential CTCF and cohesin peaks in ENCODE mouse consortium datasets.**

**A.** The x-axis indicates the number of male mouse tissues other than liver where CTCF is bound, out of 15 tissues examined by the ENCODE Consortium. A value of 15 indicates tissue-ubiquitous CTCF binding, and a value of 0 indicates liver-specific CTCF binding. The y-axis shows the proportion of male-biased peaks (blue) or female-biased peaks (red) that fall into a given bin. P-values comparing the distribution of tissue-specificity values for CTCF binding between males and females are indicated in the upper *left* corner of each plot (K-S t-test). Results show that CAC sites (upper panels) are much less tissue-specific than Lone CTCF and CNC sites (lower panels). Further, female-biased CAC peaks are less tissue-specific than male-biased CAC peaks, while male-biased Lone CTCF peaks are less liver-specific than female-biased peaks of the same class. The greater tissue ubiquity of CTCF binding for female-biased CAC peaks could be due to the fact that female-biased CTCF peaks are stronger and contain higher quality CTCF motifs, insofar as stronger peaks show greater conservation for both sex-differential and all CTCF peaks (see panel B, below); however, male-biased Lone CTCF peaks are not significantly stronger, nor do they contain higher quality motifs than the female-biased Lone CTCF peaks. The apparent difference could be due to the fact that the CTCF ChIP-seq data from the non-liver, non-reproductive tissues examined here was obtained by the ENCODE consortium from male mice [52]. Very few male-biased and female-biased CNC peaks were bound by CTCF in other any other mouse tissues (<20% of the total sex-biased liver CNC sites). This finding provides additional evidence that CNC sites are found at liver-specific *cis* regulatory elements, and that these sites do not act as insulators in other non-liver tissues (i.e., CTCF binding is lost in liver or gained in some other tissue).

**B.** The tissue specificity of neighboring genes varies significantly with the class of CTCF/cohesin binding site. Shown are Tau values, where a value close to 1 indicates the pattern of expression across mouse ENCODE RNA-seq datasets is highly tissue-specific, and where Tau values less than ∼0.3 indicate housekeeping genes. Nearest genes (within 20 kb) were defined based on distance to the TSS, and only liver-expressed genes were considered (FPKM > 1). Tau values were calculated based on the average of two replicates from all tissues except testis, using expression data generated by the ENCODE consortium. Both female-biased and male-biased CNC sites are near genes that generally are more tissue-specific than liver-expressed genes. In addition, genes near female-biased CNC sites are significantly different than genes near similarly sex-biased CACs (p=0.007; M-W test). This difference is not a reflection of the male-biased or female-biased CAC group used in the comparison, insofar as genes near female-biased CNCs are significantly more liver-specific than genes near male-biased CAC sites (p=0.0171; M-W test), while the opposite comparison for male-biased CNCs vs female-biased CACs is still not significant (p=0.07; M-W test). For these analyses, liver-expressed genes are defined by a liver expression value of FPKM > 1 (8,810 genes in total) and mapping was based on the closest TSS within 20 kb of a peak.

**C.** There is a strong association between CTCF peak strength and tissue conservation of CTCF binding, which likely explains the modestly higher tissue conservation of female-biased CTCF and cohesin peaks seen in panel A. Shown on the y-axis are reads-in-peaks normalized ChIP-seq data for all CTCF peaks, male-biased CTCF peaks, and female-biased CTCF peaks. These are grouped according to the number of non-liver ENCODE tissues with a CTCF peak, where 0 indicates a peak is liver-specific and 15 means all male mouse tissues with ENCODE datasets have CTCF bound at that position, as in panel A.

**Fig. S4 − Screenshot of TAD containing C8a/C8b, and cohesin depletion effects**

**A.** Shown is a screenshot with proposed model linking the distal male-biased enhancers and component complement genes *C8a* and *C8b* within a single TAD on mouse chromosome 4. This screenshot spans chr4:102960671-104603975. The tracks, normalization, and annotations are as described in Fig. 3.

**B.** For the proximally-regulated male-biased genes shown in Fig. 3B (*Nat8* and *Cml5*), depletion of cohesin does not significantly impact gene expression.

**Fig. S5 − 4C-seq data tracks for individual male and female mouse livers, and expression of A1bg region sex-specific lncRNAs.**

**A-D**. Shown are 4C-seq data for the same genomic regions presented in Fig. 4, but showing the individual 4C-seq data for each of 3 male liver and 3 female liver biological replicates per viewpoint. The gene tracks and sex-biased sites are as described in Fig. 4. These bed file tracks are as follows (from top to bottom): sex-biased H3K27ac peaks, sex-biased DHS, sex-biased cohesin peaks, and sex-biased CTCF peaks. Protein coding genes, sex-biased lncRNA genes, and intra-TAD loops are also shown, where present.

**A.** All six biological replicates for the *A1bg* enhancer viewpoint (chr15:60733512-60954051).

**B.** All six biological replicates for the *Gm4794* enhancer viewpoint (chr10:33418446-33680888).

**C.** All six biological replicates for the *C9* promoter viewpoint (chr15:6147917-6461799).

**D.** All six biological replicates for the *Nudt7* enhancer viewpoint (chr8:116592444-116707613).

**E.** Shown is RNA-seq expression data for *A1bg* and 12 mono-exonic lncRNAs (see Fig. 4A), obtained in six separate RNA-seq datasets from CD-1 mouse liver, and one RNA-seq dataset from C57/Bl6 mouse liver. The first two columns indicate the maximum expression (in FPKM) for male and female liver across these datasets. Following this from left to right, the columns indicate the mean expression level of each gene in female liver (FPKM values) for: Total PolyA+ unstranded RNA-seq [sequencing series G83; [9]], Total PolyA+ unstranded RNA-seq [sequencing series G85; [14]], Total PolyA+ stranded RNA-seq [sequencing series G118; [9]], Nuclear PolyA+ stranded RNA-seq [sequencing series G119; [9]], Total Ribosomal RNA-depleted stranded RNA-seq [sequencing series G118; [9]], Nuclear Ribosomal RNA-depleted stranded RNA-seq [sequencing series G119; [9]], and Total Ribosomal RNA-depleted stranded RNA-seq [from C57Bl6/J, all others CD-1; [97]]. The final columns indicate the nuclear enrichment for PolyA+ RNA datasets and for Ribosomal RNA-depleted RNA-seq datasets (linear scale). Specifically, for PolyA+ datasets, this is the FPKM value in data column 6 (“Nuclear Poly A Strnd”) divided by the FPKM value in data column 5 (“Total Poly A Strnd”). Similarly, the final column is calculated for Ribosomal RNA-Depleted Nuclear versus total RNA-seq expression (FPKM in data column 8 “Nuclear RiboM Strnd” dividied by the FPKM value in data column 7 “Total RiboM Strnd”).

**F.** Shown are the log2 male/female expression ratios for *A1bg* and the 12 mono-exonic lncRNAs (Fig. 4A) for the seven RNA-seq datasets described in panel C. Fold change is calculated by EdgeR for all datasets and the order of datasets is the same as in panel C. The final column indicates the number of datasets in which the sex differences indicated are significant (FDR < 0 .05; EdgeR).

**G.** Distal enhancer regions from panel C with bidirectional eRNA loci and YY1 binding sites indicated in relation to male-biased enhancers (chr15:6,164,877-6,184,670).

**Fig. S6 − Gel analysis/quality control of 4C-seq libraries**

**A.** Agarose gel analysis for quality control of ligated, digested, and re-ligated 4C samples. Lane (i) analyzes a sample after proximity ligation, lane (ii) shows the sample after digestion with the restriction enzyme Csp6i, and lane (iii) shows the sample after self-circularization ligation. Lane (iii) represents the final material used as input for inverse PCR with viewpoint-specific primers (Table S3A). DNA fragment sizes (in kb) are marked on the *left* of the gel.

**B.** Agarose gel analysis for quality control of final 4C-seq libraries after the inverse PCR step. A diverse mixture of PCR products is present, as indicated by a smear on the gel, with sizes primarily below ∼1 kb, which indicate a high-quality library and which allows for efficient sequencing. DNA fragment sizes (in kb) are marked on the *right* of the gel.

**Fig. S7 − High expression of Nox4 in hepatocellular carcinoma and 4C-seq biological replicates**

**A.** *Nox4* is highly up regulated in tumor relative to normal healthy tissue of mice that spontaneously develop tumors (mouse strain C3H; [95]) (p=0.0006, M-W t-test).

**B.** In human patient samples, *Nox4* is consistently up regulated in tumor tissue relative to normal controls, including in hepatocellular carcinoma (marked by thick black line). Only data for cancer types with matched primary tumor and normal tissue controls are shown, with *Nox4* showing significant up regulation in tumors for 14 of 18 matched tissue pairs. Expression datasets are from The Cancer Genome Atlas (TCGA) and were analyzed by the Tumor IMmune Estimation Resource (TIMER) webtool (https://cistrome.shinyapps.io/timer/) with default parameters [94]. The significance of comparisons between Normal and Tumor tissue was calculated by Wilcoxon test and is indicated on the chart as: 0 ≤ *** < 0.001 ≤ ** < 0.01 ≤ * < 0.05. For example, the difference between liver tumor versus normal tissue (p = 1.35E-25) is indicated as ***.

**C.** Chromatin state key for *top* browser track in Fig. 5A, which is based on chromatin states in both male and female mouse liver, which were determined for the entire genome based on a 14-state model of chromatin states developed in [8].

**D.** Male and female 4C-seq biological replicates for the *Nox4* enhancer-1 viewpoint (chr7:94248242-94726358). Tracks are as described in Fig. S5. “SB Peaks” indicate the male-biased (blue) and female-biased (pink) for the indicated factor or assay and a darker color indicates a more stringently-defined sex-biased region (cutoffs differ for each factor; see Methods). These tracks, from top to bottom, are as follows: H3K27ac ChIP-seq, DNase-seq, Rad21 ChIP-seq, and CTCF ChIP-seq and are as described in Fig. 3 and Fig. 4. A complete listing of all such peaks is provided in Tables 2D and 2E (K27ac ChIP-seq and DNase-seq) and in Tables 1D, and 1E (CTCF and cohesin ChIP-seq peaks).

**E.** Male and female 4C-seq biological replicates for the *Nox4* enhancer-2 viewpoint (chr7:94248242-94726358). Tracks are as described in Fig. S5 and in panel D of this figure.

**F.** Three separate models of regulatory domain prediction support the model that *Nox4* enhancer-1 viewpoint and enhancer-2 viewpoint (VP1, VP2) do not interact. The position of enhancer-1 viewpoint is indicated by a red arrow, and the position of enhancer-2 viewpoint is indicated by a green arrow. From *top* to *bottom*: Enhancer Promoter Units (EPUs) are domains based on the pan-tissue correlation of ChIP-seq signal at enhancer and promoters [52]; intra-TAD loops are computationally predicted based on CTCF and cohesin ChIP-seq data for male mouse liver [32]; cohesin ChIA-PET loops show cohesin-anchored interactions from mouse embryonic stem cells [27].

## References

1. Mayne BT, Bianco-Miotto T, Buckberry S, Breen J, Clifton V, Shoubridge C, Roberts CT: Large Scale Gene Expression Meta-Analysis Reveals Tissue-Specific, Sex-Biased Gene Expression in Humans. Front Genet 2016, 7:183.

2. Qu K, Zaba LC, Giresi PG, Li R, Longmire M, Kim YH, Greenleaf WJ, Chang HY: Individuality and variation of personal regulomes in primary human T cells. Cell Syst 2015, 1(1):51–61.

3. Rinn JL, Rozowsky JS, Laurenzi IJ, Petersen PH, Zou K, Zhong W, Gerstein M, Snyder M: Major molecular differences between mammalian sexes are involved in drug metabolism and renal function. Dev Cell 2004, 6(6):791–800.

4. Zhang Y, Klein K, Sugathan A, Nassery N, Dombkowski A, Zanger UM, Waxman DJ: Transcriptional profiling of human liver identifies sex-biased genes associated with polygenic dyslipidemia and coronary artery disease. PLoS One 2011, 6(8):e23506.

5. Ruggieri A, Barbati C, Malorni W: Cellular and molecular mechanisms involved in hepatocellular carcinoma gender disparity. Int J Cancer 2010, 127(3):499–504.

6. Guy J, Peters MG: Liver disease in women: the influence of gender on epidemiology, natural history, and patient outcomes. Gastroenterol Hepatol (N Y) 2013, 9(10):633–639.

7. Buzzetti E, Parikh PM, Gerussi A, Tsochatzis E: Gender differences in liver disease and the drug-dose gender gap. Pharmacol Res 2017, 120:97–108.

8. Sugathan A, Waxman DJ: Genome-wide analysis of chromatin states reveals distinct mechanisms of sex-dependent gene regulation in male and female mouse liver. Mol Cell Biol 2013, 33(18):3594–3610.

9. Lau-Corona D, Suvorov A, Waxman DJ: Feminization of Male Mouse Liver by Persistent Growth Hormone Stimulation: Activation of Sex-Biased Transcriptional Networks and Dynamic Changes in Chromatin States. Mol Cell Biol 2017, 37(19).

10. Clodfelter KH, Holloway MG, Hodor P, Park SH, Ray WJ, Waxman DJ: Sex-dependent liver gene expression is extensive and largely dependent upon signal transducer and activator of transcription 5b (STAT5b): STAT5b-dependent activation of male genes and repression of female genes revealed by microarray analysis. Mol Endocrinol 2006, 20(6):1333–1351.

11. Melia T, Hao P, Yilmaz F, Waxman DJ: Hepatic Long Intergenic Noncoding RNAs: High Promoter Conservation and Dynamic, Sex-Dependent Transcriptional Regulation by Growth Hormone. Mol Cell Biol 2016, 36(1):50–69.

12. Melia T, Waxman DJ: Sex-Biased lncRNAs Inversely Correlate With Sex-Opposite Gene Coexpression Networks in Diversity Outbred Mouse Liver. Endocrinology 2019, 160(5):989–1007.

13. Hao P, Waxman DJ: Functional Roles of Sex-Biased, Growth Hormone-Regulated MicroRNAs miR-1948 and miR-802 in Young Adult Mouse Liver. Endocrinology 2018, 159(3):1377-1392.

14. Connerney J, Lau-Corona D, Rampersaud A, Waxman DJ: Activation of Male Liver Chromatin Accessibility and STAT5-Dependent Gene Transcription by Plasma Growth Hormone Pulses. Endocrinology 2017, 158(5):1386–1405.

15. Ling G, Sugathan A, Mazor T, Fraenkel E, Waxman DJ: Unbiased, genome-wide in vivo mapping of transcriptional regulatory elements reveals sex differences in chromatin structure associated with sex-specific liver gene expression. Mol Cell Biol 2010, 30(23):5531–5544.

16. Phillips JE, Corces VG: CTCF: master weaver of the genome. Cell 2009, 137(7):1194–1211.

17. Ghirlando R, Felsenfeld G: CTCF: making the right connections. Genes Dev 2016, 30(8):881–891.

18. Sanborn AL, Rao SS, Huang SC, Durand NC, Huntley MH, Jewett AI, Bochkov ID, Chinnappan D, Cutkosky A, Li J et al: Chromatin extrusion explains key features of loop and domain formation in wild-type and engineered genomes. Proceedings of the National Academy of Sciences of the United States of America 2015, 112(47):E6456–6465.

19. Davidson IF, Goetz D, Zaczek MP, Molodtsov MI, Huis In’t Veld PJ, Weissmann F, Litos G, Cisneros DA, Ocampo-Hafalla M, Ladurner R et al: Rapid movement and transcriptional re-localization of human cohesin on DNA. EMBO J 2016, 35(24):2671–2685.

20. Fedoriw AM, Stein P, Svoboda P, Schultz RM, Bartolomei MS: Transgenic RNAi reveals essential function for CTCF in H19 gene imprinting. Science 2004, 303(5655):238-240.

21. Xu H, Balakrishnan K, Malaterre J, Beasley M, Yan Y, Essers J, Appeldoorn E, Tomaszewski JM, Vazquez M, Verschoor S et al: Rad21-cohesin haploinsufficiency impedes DNA repair and enhances gastrointestinal radiosensitivity in mice. PLoS One 2010, 5(8):e12112.

22. Schwarzer W, Abdennur N, Goloborodko A, Pekowska A, Fudenberg G, Loe-Mie Y, Fonseca NA, Huber W, Haering CH, Mirny L et al: Two independent modes of chromatin organization revealed by cohesin removal. Nature 2017, 551(7678):51-56.

23. Rao SSP, Huang SC, St Hilaire BG, Engreitz JM, Perez EM, Kieffer-Kwon KR, Sanborn AL, Johnstone SE, Bascom GD, Bochkov ID et al: Cohesin Loss Eliminates All Loop Domains. Cell 2017, 171(2):305-+.

24. Nora EP, Goloborodko A, Valton AL, Gibcus JH, Uebersohn A, Abdennur N, Dekker J, Mirny LA, Bruneau BG: Targeted Degradation of CTCF Decouples Local Insulation of Chromosome Domains from Genomic Compartmentalization. Cell 2017, 169(5):930–944 e922.

25. Ciosk R, Shirayama M, Shevchenko A, Tanaka T, Toth A, Shevchenko A, Nasmyth K: Cohesin’s binding to chromosomes depends on a separate complex consisting of Scc2 and Scc4 proteins. Mol Cell 2000, 5(2):243–254.

26. Kagey MH, Newman JJ, Bilodeau S, Zhan Y, Orlando DA, van Berkum NL, Ebmeier CC, Goossens J, Rahl PB, Levine SS et al: Mediator and cohesin connect gene expression and chromatin architecture. Nature 2010, 467(7314):430-435.

27. Dowen JM, Fan ZP, Hnisz D, Ren G, Abraham BJ, Zhang LN, Weintraub AS, Schujiers J, Lee TI, Zhao K et al: Control of cell identity genes occurs in insulated neighborhoods in mammalian chromosomes. Cell 2014, 159(2):374–387.

28. Phillips-Cremins JE, Sauria ME, Sanyal A, Gerasimova TI, Lajoie BR, Bell JS, Ong CT, Hookway TA, Guo C, Sun Y et al: Architectural protein subclasses shape 3D organization of genomes during lineage commitment. Cell 2013, 153(6):1281–1295.

29. Faure AJ, Schmidt D, Watt S, Schwalie PC, Wilson MD, Xu H, Ramsay RG, Odom DT, Flicek P: Cohesin regulates tissue-specific expression by stabilizing highly occupied cis-regulatory modules. Genome Res 2012, 22(11):2163–2175.

30. Schmidt D, Schwalie PC, Ross-Innes CS, Hurtado A, Brown GD, Carroll JS, Flicek P, Odom DT: A CTCF-independent role for cohesin in tissue-specific transcription. Genome Res 2010, 20(5):578–588.

31. Dixon JR, Selvaraj S, Yue F, Kim A, Li Y, Shen Y, Hu M, Liu JS, Ren B: Topological domains in mammalian genomes identified by analysis of chromatin interactions. Nature 2012, 485(7398):376-380.

32. Matthews BJ, Waxman DJ: Computational prediction of CTCF/cohesin-based intra-TAD loops that insulate chromatin contacts and gene expression in mouse liver. Elife 2018, 7.

33. Rao SS, Huntley MH, Durand NC, Stamenova EK, Bochkov ID, Robinson JT, Sanborn AL, Machol I, Omer AD, Lander ES et al: A 3D map of the human genome at kilobase resolution reveals principles of chromatin looping. Cell 2014, 159(7):1665–1680.

34. Dixon JR, Jung I, Selvaraj S, Shen Y, Antosiewicz-Bourget JE, Lee AY, Ye Z, Kim A, Rajagopal N, Xie W et al: Chromatin architecture reorganization during stem cell differentiation. Nature 2015, 518(7539):331-336.

35. Vietri Rudan M, Barrington C, Henderson S, Ernst C, Odom DT, Tanay A, Hadjur S: Comparative Hi-C reveals that CTCF underlies evolution of chromosomal domain architecture. Cell Rep 2015, 10(8):1297–1309.

36. Ren G, Jin W, Cui K, Rodrigez J, Hu G, Zhang Z, Larson DR, Zhao K: CTCF-Mediated Enhancer-Promoter Interaction Is a Critical Regulator of Cell-to-Cell Variation of Gene Expression. Mol Cell 2017, 67(6):1049–1058 e1046.

37. Splinter E, Heath H, Kooren J, Palstra RJ, Klous P, Grosveld F, Galjart N, de Laat W: CTCF mediates long-range chromatin looping and local histone modification in the beta-globin locus. Genes Dev 2006, 20(17):2349–2354.

38. Huang J, Li K, Cai W, Liu X, Zhang Y, Orkin SH, Xu J, Yuan GC: Dissecting super-enhancer hierarchy based on chromatin interactions. Nat Commun 2018, 9(1):943.

39. Donohoe ME, Zhang LF, Xu N, Shi Y, Lee JT: Identification of a Ctcf cofactor, Yy1, for the X chromosome binary switch. Mol Cell 2007, 25(1):43-56.

40. Pugacheva EM, Rivero-Hinojosa S, Espinoza CA, Mendez-Catala CF, Kang S, Suzuki T, Kosaka-Suzuki N, Robinson S, Nagarajan V, Ye Z et al: Comparative analyses of CTCF and BORIS occupancies uncover two distinct classes of CTCF binding genomic regions. Genome Biol 2015, 16(1):161.

41. Park HJ, Li J, Hannah R, Biddie S, Leal-Cervantes AI, Kirschner K, Flores Santa Cruz D, Sexl V, Gottgens B, Green AR: Cytokine-induced megakaryocytic differentiation is regulated by genome-wide loss of a uSTAT transcriptional program. EMBO J 2016, 35(6):580–594.

42. Guo Y, Xu Q, Canzio D, Shou J, Li J, Gorkin DU, Jung I, Wu H, Zhai Y, Tang Y et al: CRISPR Inversion of CTCF Sites Alters Genome Topology and Enhancer/Promoter Function. Cell 2015, 162(4):900–910.

43. Oti M, Falck J, Huynen MA, Zhou H: CTCF-mediated chromatin loops enclose inducible gene regulatory domains. BMC Genomics 2016, 17:252.

44. Whalen S, Truty RM, Pollard KS: Enhancer–promoter interactions are encoded by complex genomic signatures on looping chromatin. Nature Genetics 2016, 48(5):488.

45. Yang Y, Zhang R, Singh S, Ma J: Exploiting sequence-based features for predicting enhancer-promoter interactions. Bioinformatics 2017, 33(14):i252–i260.

46. Dekker J, Rippe K, Dekker M, Kleckner N: Capturing chromosome conformation. Science 2002, 295(5558):1306-1311.

47. van de Werken HJ, Landan G, Holwerda SJ, Hoichman M, Klous P, Chachik R, Splinter E, Valdes-Quezada C, Oz Y, Bouwman BA et al: Robust 4C-seq data analysis to screen for regulatory DNA interactions. Nat Methods 2012, 9(10):969–972.

48. Hark AT, Schoenherr CJ, Katz DJ, Ingram RS, Levorse JM, Tilghman SM: CTCF mediates methylation-sensitive enhancer-blocking activity at the H19/Igf2 locus. Nature 2000, 405(6785):486-489.

49. Renda M, Baglivo I, Burgess-Beusse B, Esposito S, Fattorusso R, Felsenfeld G, Pedone PV: Critical DNA binding interactions of the insulator protein CTCF - A small number of zinc fingers mediate strong binding, and a single finger-DNA interaction controls binding at imprinted loci. Journal of Biological Chemistry 2007, 282(46):33336–33345.

50. Reizel Y, Spiro A, Sabag O, Skversky Y, Hecht M, Keshet I, Berman BP, Cedar H: Gender-specific postnatal demethylation and establishment of epigenetic memory. Genes Dev 2015, 29(9):923–933.

51. Seitan VC, Faure AJ, Zhan Y, McCord RP, Lajoie BR, Ing-Simmons E, Lenhard B, Giorgetti L, Heard E, Fisher AG et al: Cohesin-based chromatin interactions enable regulated gene expression within preexisting architectural compartments. Genome Res 2013, 23(12):2066–2077.

52. Shen Y, Yue F, McCleary DF, Ye Z, Edsall L, Kuan S, Wagner U, Dixon J, Lee L, Lobanenkov VV et al: A map of the cis-regulatory sequences in the mouse genome. Nature 2012, 488(7409):116-120.

53. Lin S, Lin Y, Nery JR, Urich MA, Breschi A, Davis CA, Dobin A, Zaleski C, Beer MA, Chapman WC et al: Comparison of the transcriptional landscapes between human and mouse tissues. Proceedings of the National Academy of Sciences of the United States of America 2014, 111(48):17224–17229.

54. Haller F, Bieg M, Will R, Korner C, Weichenhan D, Bott A, Ishaque N, Lutsik P, Moskalev EA, Mueller SK et al: Enhancer hijacking activates oncogenic transcription factor NR4A3 in acinic cell carcinomas of the salivary glands. Nat Commun 2019, 10(1):368.

55. Weintraub AS, Li CH, Zamudio AV, Sigova AA, Hannett NM, Day DS, Abraham BJ, Cohen MA, Nabet B, Buckley DL: YY1 is a structural regulator of enhancer-promoter loops. Cell 2017, 171(7):1573–1588. e1528.

56. Engreitz JM, Pandya-Jones A, McDonel P, Shishkin A, Sirokman K, Surka C, Kadri S, Xing J, Goren A, Lander ES et al: The Xist lncRNA exploits three-dimensional genome architecture to spread across the X chromosome. Science 2013, 341(6147):1237973.

57. Hacisuleyman E, Goff LA, Trapnell C, Williams A, Henao-Mejia J, Sun L, McClanahan P, Hendrickson DG, Sauvageau M, Kelley DR et al: Topological organization of multichromosomal regions by the long intergenic noncoding RNA Firre. Nat Struct Mol Biol 2014, 21(2):198–206.

58. Yin Y, Yan P, Lu J, Song G, Zhu Y, Li Z, Zhao Y, Shen B, Huang X, Zhu H et al: Opposing Roles for the lncRNA Haunt and Its Genomic Locus in Regulating HOXA Gene Activation during Embryonic Stem Cell Differentiation. Cell Stem Cell 2015, 16(5):504–516.

59. Kung JT, Kesner B, An JY, Ahn JY, Cifuentes-Rojas C, Colognori D, Jeon Y, Szanto A, del Rosario BC, Pinter SF et al: Locus-specific targeting to the X chromosome revealed by the RNA interactome of CTCF. Mol Cell 2015, 57(2):361–375.

60. Schwartzman O, Mukamel Z, Oded-Elkayam N, Olivares-Chauvet P, Lubling Y, Landan G, Izraeli S, Tanay A: UMI-4C for quantitative and targeted chromosomal contact profiling. Nat Methods 2016, 13(8):685–691.

61. Borrie MS, Campor JS, Joshi H, Gartenberg MR: Binding, sliding, and function of cohesin during transcriptional activation. Proceedings of the National Academy of Sciences of the United States of America 2017, 114(7):E1062–E1071.

62. Busslinger GA, Stocsits RR, van der Lelij P, Axelsson E, Tedeschi A, Galjart N, Peters JM: Cohesin is positioned in mammalian genomes by transcription, CTCF and Wapl. Nature 2017, 544(7651):503-507.

63. Liang S, Kisseleva T, Brenner DA: The Role of NADPH Oxidases (NOXs) in Liver Fibrosis and the Activation of Myofibroblasts. Frontiers in physiology 2016, 7:17.

64. Sun Q, Zhang W, Zhong W, Sun X, Zhou Z: Pharmacological inhibition of NOX4 ameliorates alcohol-induced liver injury in mice through improving oxidative stress and mitochondrial function. Biochimica et biophysica acta General subjects 2017, 1861(1 Pt A):2912–2921.

65. Li W, Notani D, Ma Q, Tanasa B, Nunez E, Chen AY, Merkurjev D, Zhang J, Ohgi K, Song X et al: Functional roles of enhancer RNAs for oestrogen-dependent transcriptional activation. Nature 2013, 498(7455):516-520.

66. Sigova AA, Abraham BJ, Ji X, Molinie B, Hannett NM, Guo YE, Jangi M, Giallourakis CC, Sharp PA, Young RA: Transcription factor trapping by RNA in gene regulatory elements. Science 2015, 350(6263):978-981.

67. Fang B, Everett LJ, Jager J, Briggs E, Armour SM, Feng D, Roy A, Gerhart-Hines Z, Sun Z, Lazar MA: Circadian enhancers coordinate multiple phases of rhythmic gene transcription in vivo. Cell 2014, 159(5):1140–1152.

68. Schwalie PC, Ward MC, Cain CE, Faure AJ, Gilad Y, Odom DT, Flicek P: Co-binding by YY1 identifies the transcriptionally active, highly conserved set of CTCF-bound regions in primate genomes. Genome Biol 2013, 14(12):R148.

69. Keleher MR, Zaidi R, Hicks L, Shah S, Xing X, Li D, Wang T, Cheverud JM: A high-fat diet alters genome-wide DNA methylation and gene expression in SM/J mice. BMC Genomics 2018, 19(1):888.

70. Lodato NJ, Melia T, Rampersaud A, Waxman DJ: Sex-Differential Responses of Tumor Promotion-Associated Genes and Dysregulation of Novel Long Noncoding RNAs in Constitutive Androstane Receptor-Activated Mouse Liver. Toxicological sciences: an official journal of the Society of Toxicology 2017, 159(1):25–41.

71. Lodato NJ, Rampersaud A, Waxman DJ: Impact of CAR Agonist Ligand TCPOBOP on Mouse Liver Chromatin Accessibility. Toxicological sciences: an official journal of the Society of Toxicology 2018, 164(1):115–128.

72. Leung A, Trac C, Du J, Natarajan R, Schones DE: Persistent Chromatin Modifications Induced by High Fat Diet. J Biol Chem 2016, 291(20):10446–10455.

73. Marques AC, Hughes J, Graham B, Kowalczyk MS, Higgs DR, Ponting CP: Chromatin signatures at transcriptional start sites separate two equally populated yet distinct classes of intergenic long noncoding RNAs. Genome Biology 2013, 14(11):R131.

74. Gil N, Ulitsky I: Production of Spliced Long Noncoding RNAs Specifies Regions with Increased Enhancer Activity. Cell Syst 2018, 7(5):537–547 e533.

75. Lieberman-Aiden E, van Berkum NL, Williams L, Imakaev M, Ragoczy T, Telling A, Amit I, Lajoie BR, Sabo PJ, Dorschner MO et al: Comprehensive mapping of long-range interactions reveals folding principles of the human genome. Science 2009, 326(5950):289-293.

76. Conforto TL, Waxman DJ: Sex-specific mouse liver gene expression: genome-wide analysis of developmental changes from pre-pubertal period to young adulthood. Biology of sex differences 2012, 3:9.

77. Lowe R, Gemma C, Rakyan VK, Holland ML: Sexually dimorphic gene expression emerges with embryonic genome activation and is dynamic throughout development. BMC Genomics 2015, 16:295.

78. Wauthier V, Sugathan A, Meyer RD, Dombkowski AA, Waxman DJ: Intrinsic sex differences in the early growth hormone responsiveness of sex-specific genes in mouse liver. Mol Endocrinol 2010, 24(3):667–678.

79. Zhang Y, Laz EV, Waxman DJ: Dynamic, sex-differential STAT5 and BCL6 binding to sex-biased, growth hormone-regulated genes in adult mouse liver. Mol Cell Biol 2012, 32(4):880–896.

80. Conforto TL, Steinhardt GF, Waxman DJ: Cross Talk Between GH-Regulated Transcription Factors HNF6 and CUX2 in Adult Mouse Liver. Molecular Endocrinology 2015, 29(9):1286–1302.

81. Wang H, Maurano MT, Qu H, Varley KE, Gertz J, Pauli F, Lee K, Canfield T, Weaver M, Sandstrom R et al: Widespread plasticity in CTCF occupancy linked to DNA methylation. Genome Res 2012, 22(9):1680–1688.

82. Takasugi M, Hayakawa K, Arai D, Shiota K: Age- and sex-dependent DNA hypomethylation controlled by growth hormone in mouse liver. Mech Ageing Dev 2013, 134(7-8):331–337.

83. Xu Y, Guo W, Li P, Zhang Y, Zhao M, Fan Z, Zhao Z, Yan J: Long-Range Chromosome Interactions Mediated by Cohesin Shape Circadian Gene Expression. PLoS Genet 2016, 12(5):e1005992.

84. Yeung J, Mermet J, Jouffe C, Marquis J, Charpagne A, Gachon F, Naef F: Transcription factor activity rhythms and tissue-specific chromatin interactions explain circadian gene expression across organs. Genome Res 2018, 28(2):182–191.

85. Mermet J, Yeung J, Hurni C, Mauvoisin D, Gustafson K, Jouffe C, Nicolas D, Emmenegger Y, Gobet C, Franken P et al: Clock-dependent chromatin topology modulates circadian transcription and behavior. Genes Dev 2018, 32(5-6):347–358.

86. Phanstiel DH, Van Bortle K, Spacek D, Hess GT, Shamim MS, Machol I, Love MI, Aiden EL, Bassik MC, Snyder MP: Static and Dynamic DNA Loops form AP-1-Bound Activation Hubs during Macrophage Development. Mol Cell 2017, 67(6):1037–1048 e1036.

87. Banerjee S, Das RK, Shapiro BH: Feminization imprinted by developmental growth hormone. Molecular and cellular endocrinology 2019, 479:27–38.

88. Landt SG, Marinov GK, Kundaje A, Kheradpour P, Pauli F, Batzoglou S, Bernstein BE, Bickel P, Brown JB, Cayting P et al: ChIP-seq guidelines and practices of the ENCODE and modENCODE consortia. Genome Res 2012, 22(9):1813–1831.

89. Rampersaud A, Lodato NJ, Shin A, Waxman DJ: Widespread epigenetic changes to the enhancer landscape of mouse liver induced by a specific xenobiotic agonist ligand of the nuclear receptor CAR. Toxicological sciences: an official journal of the Society of Toxicology 2019.

90. Shen L, Shao NY, Liu X, Maze I, Feng J, Nestler EJ: diffReps: detecting differential chromatin modification sites from ChIP-seq data with biological replicates. PLoS One 2013, 8(6):e65598.

91. Shao Z, Zhang Y, Yuan GC, Orkin SH, Waxman DJ: MAnorm: a robust model for quantitative comparison of ChIP-Seq data sets. Genome Biol 2012, 13(3):R16.

92. Raviram R, Rocha PP, Muller CL, Miraldi ER, Badri S, Fu Y, Swanzey E, Proudhon C, Snetkova V, Bonneau R et al: 4C-ker: A Method to Reproducibly Identify Genome-Wide Interactions Captured by 4C-Seq Experiments. PLoS Comput Biol 2016, 12(3):e1004780.

93. Servant N, Varoquaux N, Lajoie BR, Viara E, Chen CJ, Vert JP, Heard E, Dekker J, Barillot E: HiC-Pro: an optimized and flexible pipeline for Hi-C data processing. Genome Biol 2015, 16(1):259.

94. Li T, Fan J, Wang B, Traugh N, Chen Q, Liu JS, Li B, Liu XS: TIMER: A Web Server ford Comprehensive Analysis of Tumor-Infiltrating Immune Cells. Cancer Res 2017, 77(21):e108–e110.

95. Aitken SJ, Ibarra-Soria X, Kentepozidou E, Flicek P, Feig C, Marioni JC, Odom DT: CTCF maintains regulatory homeostasis of cancer pathways. Genome Biol 2018, 19(1):106.

96. Lau-Corona D, Bae WK, Hennighausen L, Waxman DJ: Sex-biased genetic programs in liver metabolism and liver fibrosis are controlled by EZH1 and EZH2. bioRxiv 2019:577056.

97. Lowe R, Gemma C, Rakyan VK, Holland ML: Sexually dimorphic gene expression emerges with embryonic genome activation and is dynamic throughout development. BMC Genomics 2015, 16(1):295.

