## Supplemental Figures for "Impact of 3-dimensional genome organization, guided by cohesin and CTCF looping, on sex-biased chromatin interactions and gene expression in mouse liver"

Fig. S1

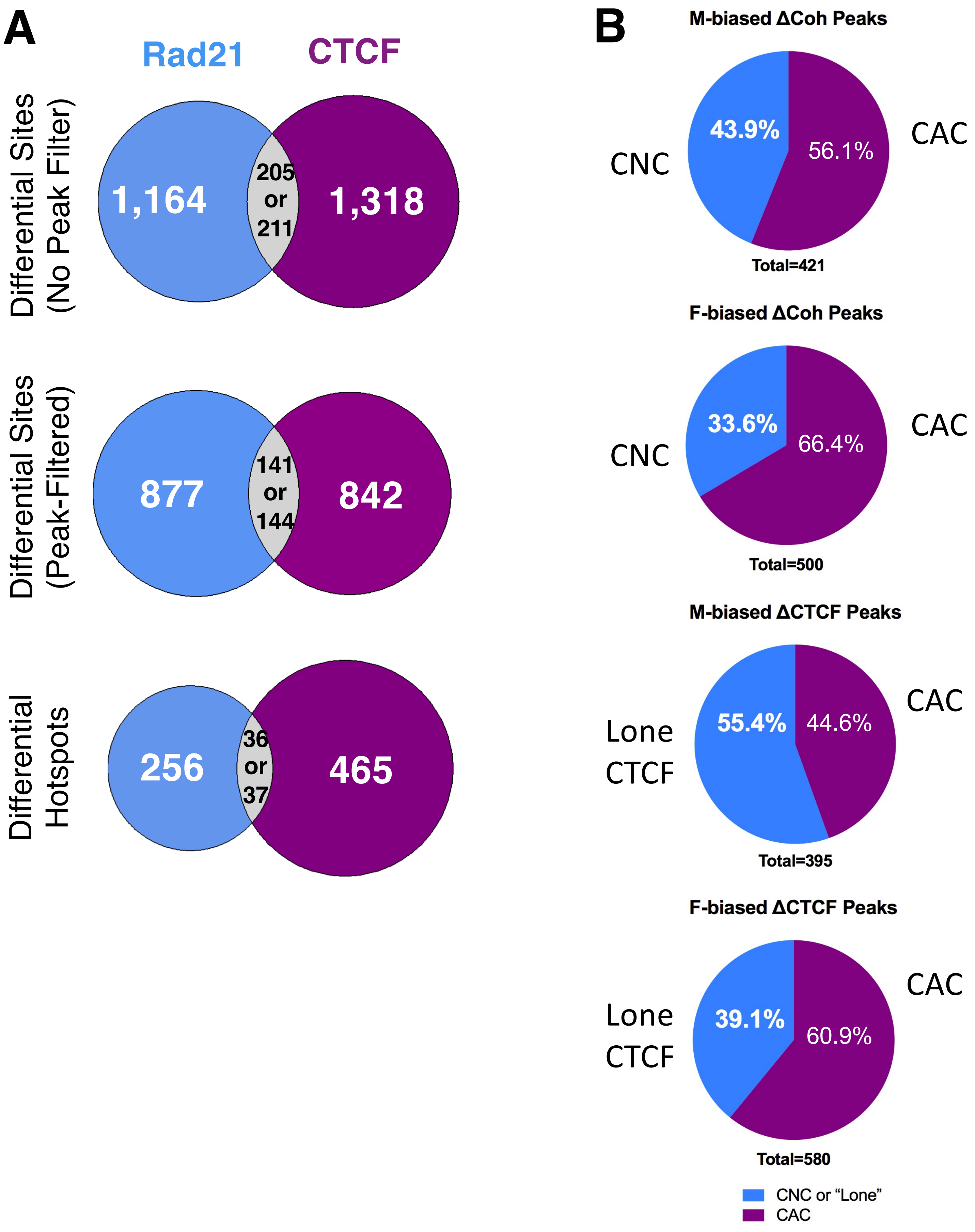

Fig. S2ABC

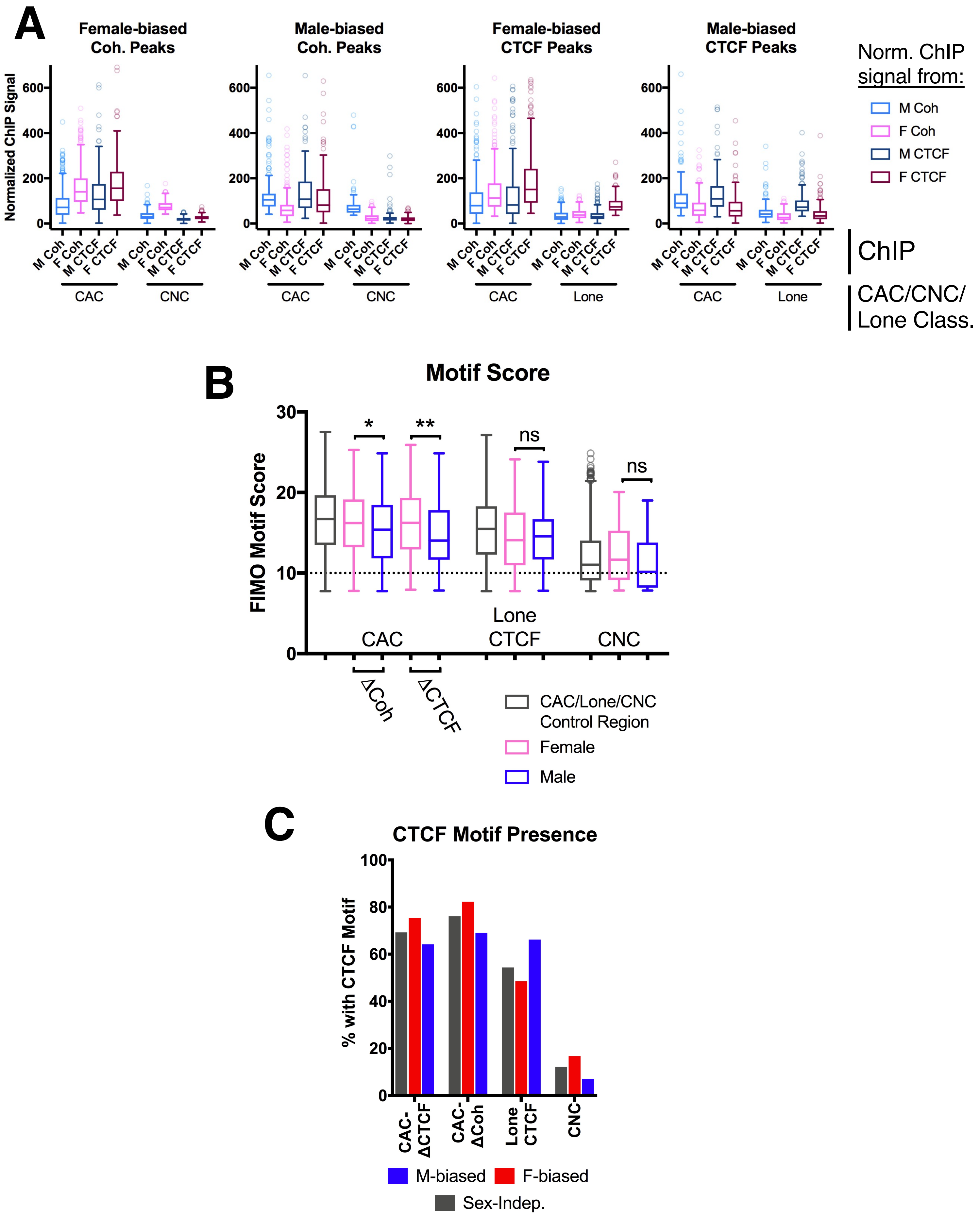

A
Female-biased Coh. Peaks
Male-biased Coh. Peaks
Female-biased CTCF Peaks
Male-biased CTCF Peaks
Normalized ChIP Signal
M Coh
F Coh
M CTCF
F CTCF
CAC
CNC
Lone
Norm. ChIP signal from:
ChIP
CAC/CNC/Lone Class.
B
Motif Score
FIMO Motif Score
ns
CAC
Lone CTCF
CNC
ΔCoh
ΔCTCF
CAC/Lone/CNC Control Region
Female
Male
C
CTCF Motif Presence
% with CTCF Motif
CAC-ΔCTCF
CAC-ΔCoh
Lone CTCF
CNC
M-biased
F-biased
Sex-Indep.

Fig. S2DE

### D Female-biased intra-TAD containing 12 sex-biased genes

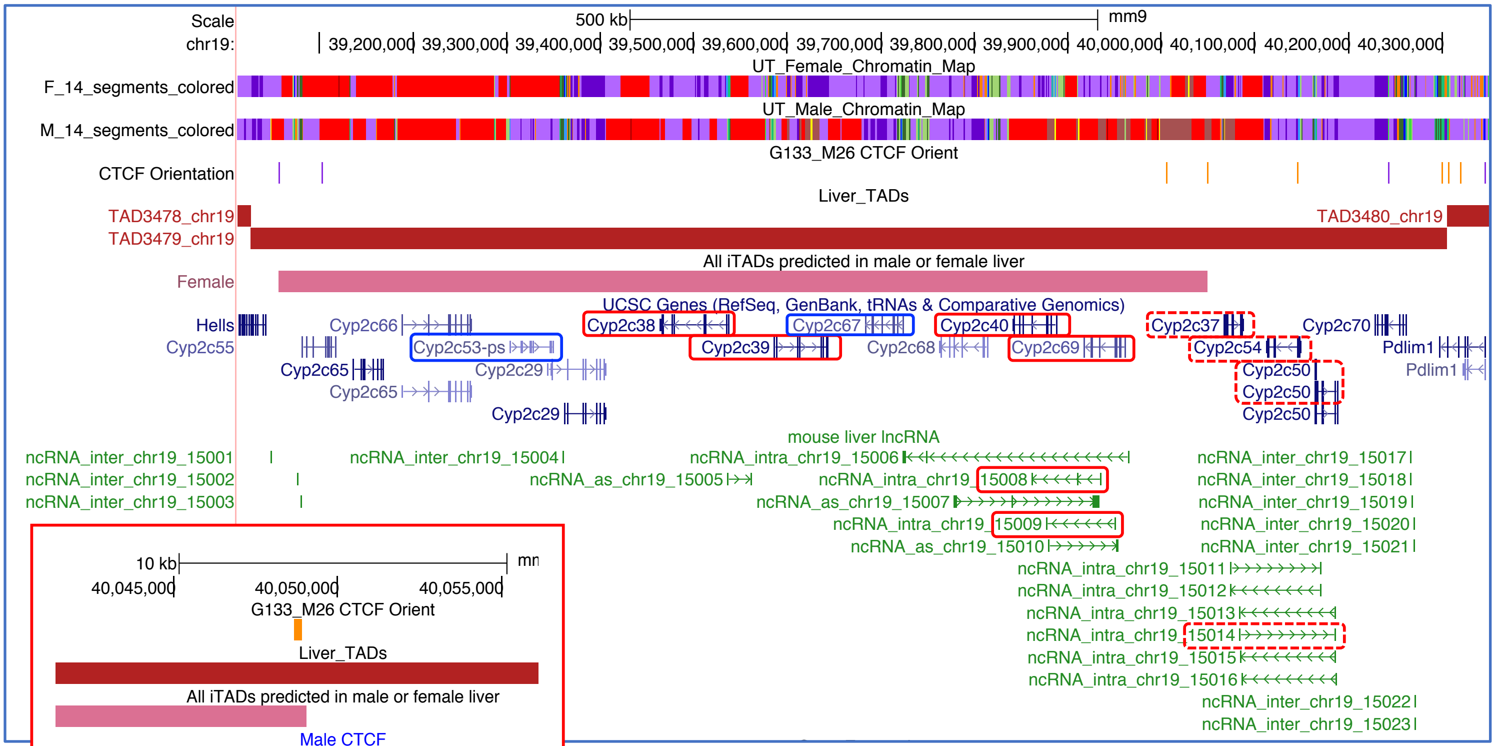


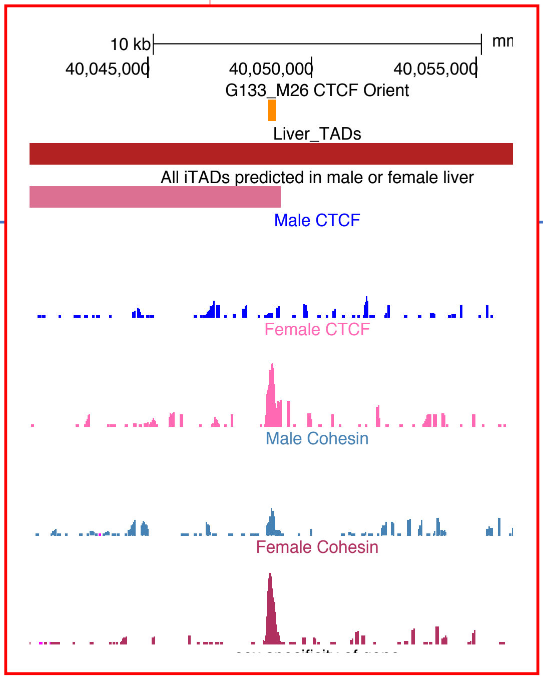


Red rectangles: female-biased gene
Blue rectangles: male-biased gene
*Dashed rectangles*: sex-biased genes in the TAD but outside the intra-TAD.

Zoom-in view of 3'-end of intra-TAD showing female-biased binding of both CTCT and cohesin (normalized ChIP-seq read wig file).

# E

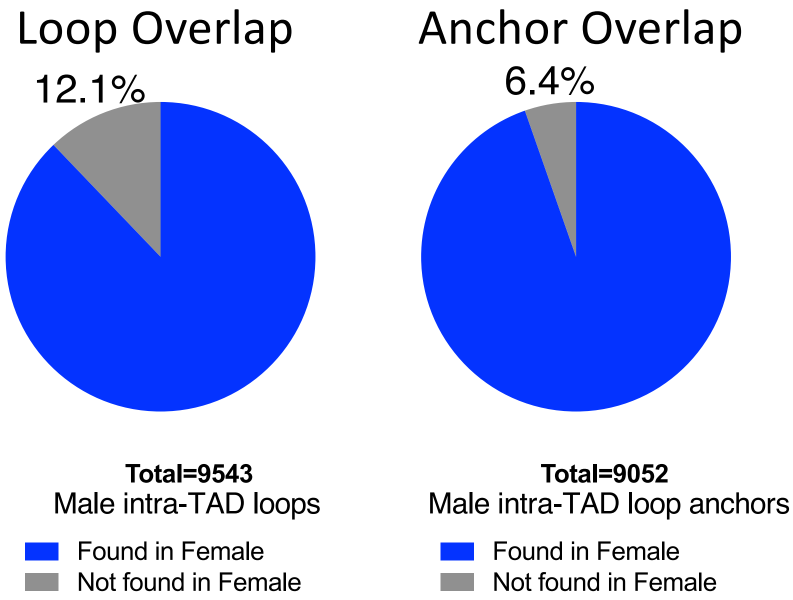

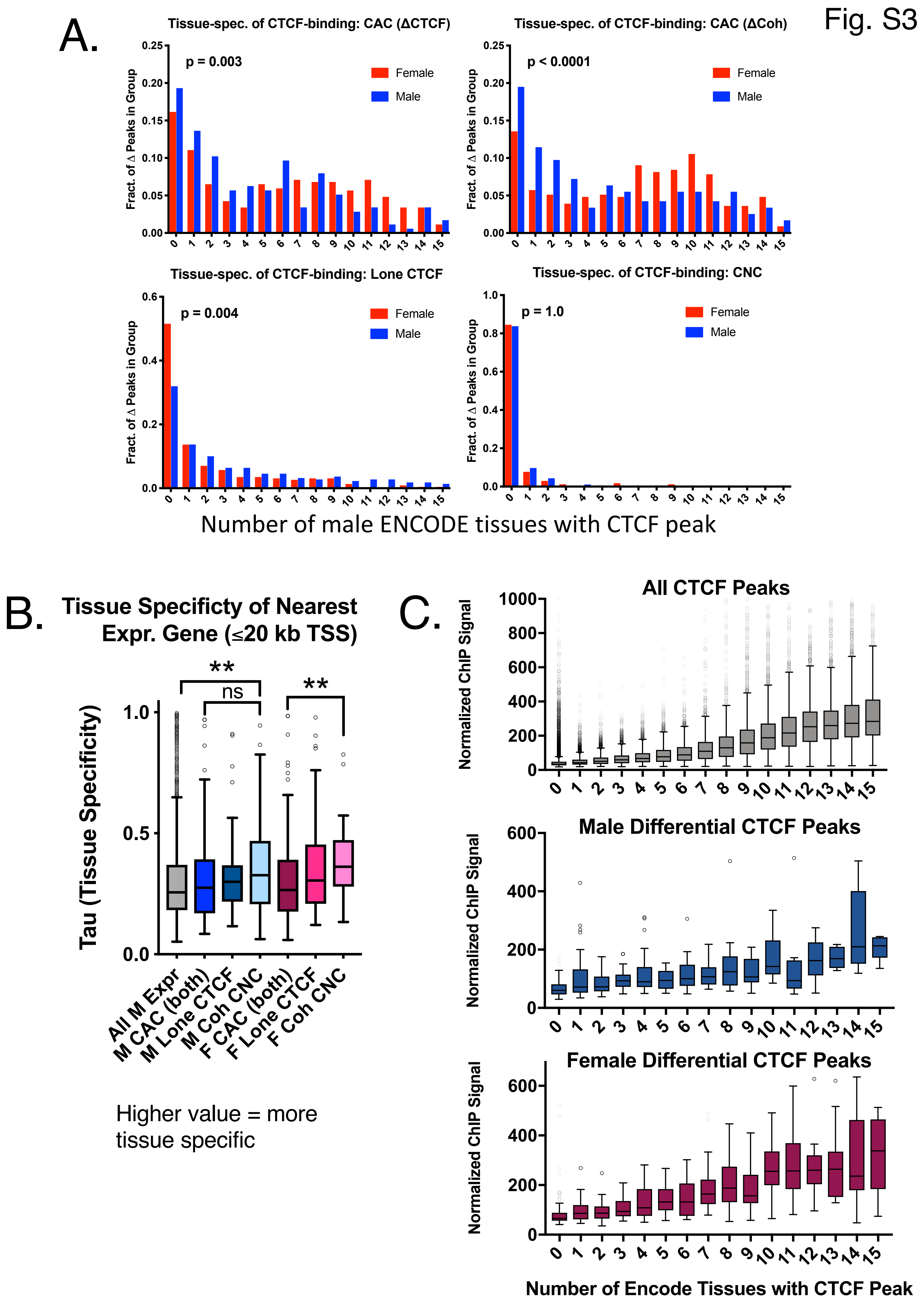
Fig. S3
A.
Tissue-spec. of CTCF-binding: CAC (ΔCTCF)
p = 0.003
Female
Male
Fract. of Δ Peaks in Group
Tissue-spec. of CTCF-binding: CAC (ΔCoh)
p < 0.0001
Tissue-spec. of CTCF-binding: Lone CTCF
p = 0.004
Tissue-spec. of CTCF-binding: CNC
p = 1.0
Number of male ENCODE tissues with CTCF peak
B.
Tissue Specificty of Nearest Expr. Gene (≤20 kb TSS)
Tau (Tissue Specificity)
**
ns
**
All M Expr
M CAC (both)
M Lone CTCF
M Coh CNC
F CAC (both)
F Lone CTCF
F Coh CNC
Higher value = more tissue specific
C.
All CTCF Peaks
Male Differential CTCF Peaks
Female Differential CTCF Peaks
Normalized ChIP Signal
Number of Encode Tissues with CTCF Peak

Fig. S4

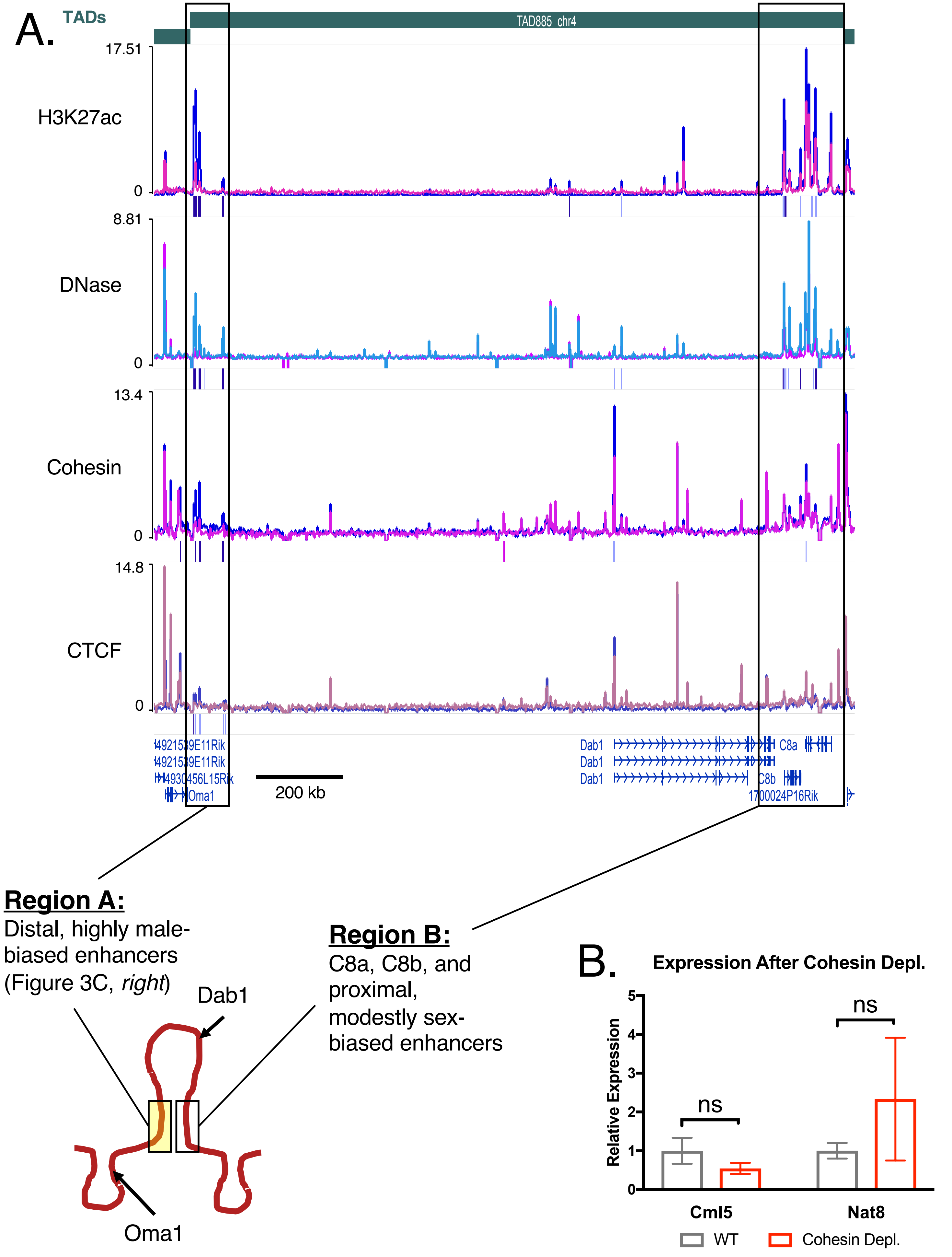
A.
TADs
TAD885_chr4
17.51
H3K27ac
0
8.81
DNase
0
13.4
Cohesin
0
14.8
CTCF
0
4921539E11Rik
4921539E11Rik
4930456L15Rik
Oma1
200 kb
Dab1
Dab1
Dab1
C8a
C8b
1700024P16Rik
Region A:
Distal, highly male-biased enhancers (Figure 3C, right)
Region B:
C8a, C8b, and proximal, modestly sex-biased enhancers
Dab1
Oma1
B.
Expression After Cohesin Depl.
Relative Expression
0
1
2
3
4
5
ns
ns
Cml5
Nat8
WT
Cohesin Depl.

Fig. S5A-S5D

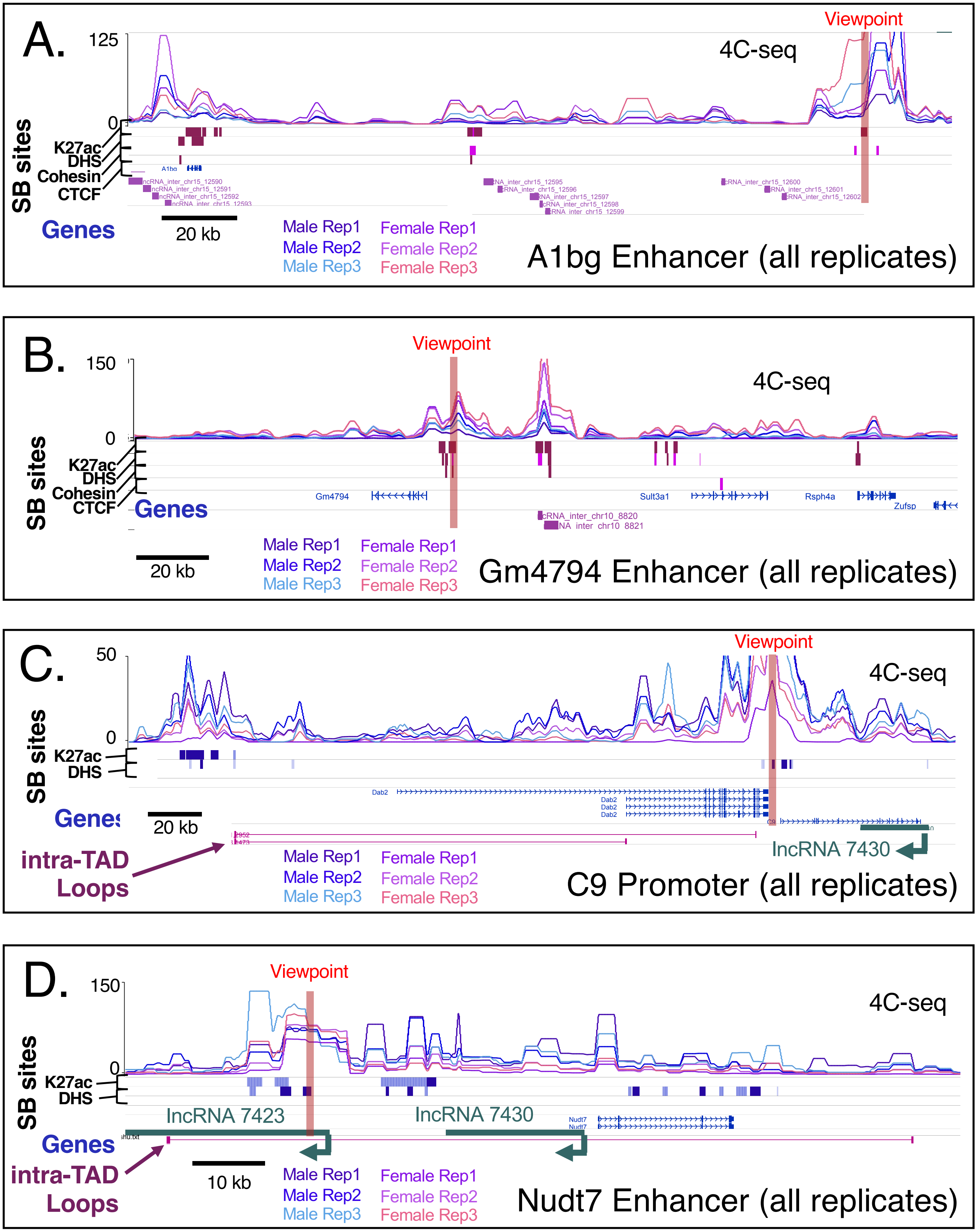

A.
Viewpoint
4C-seq
125
0
SB sites
K27ac
DHS
Cohesin
CTCF
Genes
20 kb
Male Rep1
Male Rep2
Male Rep3
Female Rep1
Female Rep2
Female Rep3
A1bg Enhancer (all replicates)
B.
Viewpoint
150
4C-seq
0
SB sites
K27ac
DHS
Cohesin
CTCF
Genes
Gm4794
Sult3a1
Rsph4a
Zufsp
20 kb
Male Rep1
Male Rep2
Male Rep3
Female Rep1
Female Rep2
Female Rep3
Gm4794 Enhancer (all replicates)
C.
50
Viewpoint
4C-seq
0
SB sites
K27ac
DHS
Dab2
Genes
20 kb
intra-TAD Loops
lncRNA 7430
Male Rep1
Male Rep2
Male Rep3
Female Rep1
Female Rep2
Female Rep3
C9 Promoter (all replicates)
D.
Viewpoint
150
4C-seq
0
SB sites
K27ac
DHS
lncRNA 7423
lncRNA 7430
Nudt7
Genes
intra-TAD Loops
10 kb
Male Rep1
Male Rep2
Male Rep3
Female Rep1
Female Rep2
Female Rep3
Nudt7 Enhancer (all replicates)

### Fig. S5E-S5G

#### E. Expression Across Datasets (F FPKM)

| | Max FPKM Male | Max FPKM Female | Total PolyA Unstr F (G83) | PolyA Unstr F (G85) | Total PolyA Strnd F | Nuclear PolyA Strnd F | Total RiboM Strnd F | Nuclear RiboM Strnd F | B6 Total RiboM Strnd F | Nuclear Enrichment: Nuclear Enrichment (PolyA) | Nuclear Enrichment: Nuclear Enrichment (RiboM) |
|---|---|---|---|---|---|---|---|---|---|---|---|
| ncRNA_inter_chr15_12590 | 0.01 | 2.71 | 0.62 | 1.30 | 0.54 | 2.57 | 0.67 | 2.71 | 0.13 | 4.8 | 4.1 |
| ncRNA_inter_chr15_12591 | 0.01 | 2.11 | 0.35 | 0.97 | 0.50 | 1.15 | 0.53 | 2.11 | 0.18 | 2.3 | 4.0 |
| ncRNA_inter_chr15_12592 | 0 | 0.99 | 0.25 | 0.35 | 0.06 | 0.22 | 0.29 | 0.99 | 0.07 | 3.6 | 3.5 |
| ncRNA_inter_chr15_12593 | 0 | 1.80 | 0.06 | 0.19 | 0.09 | 1.06 | 0.31 | 1.80 | 0.12 | 11.4 | 5.9 |
| ncRNA_inter_chr15_12595 | 0.01 | 1.13 | 0.01 | 0.01 | 0.01 | 0.33 | 0.23 | 1.13 | 0.03 | 38.8 | 5.0 |
| ncRNA_inter_chr15_12596 | 0.02 | 1.77 | 0.02 | 0 | 0.03 | 0.24 | 0.15 | 1.77 | 0.07 | 9.3 | 11.8 |
| ncRNA_inter_chr15_12597 | 0 | 0.91 | 0.01 | 0.03 | 0.02 | 0.34 | 0.09 | 0.91 | 0.04 | 15.0 | 9.9 |
| ncRNA_inter_chr15_12598 | 0 | 0.87 | 0.06 | 0.00 | 0.07 | 0.68 | 0.11 | 0.87 | 0.02 | 9.2 | 8.1 |
| ncRNA_inter_chr15_12599 | 0.02 | 0.81 | 0.02 | 0.04 | 0.03 | 0.81 | 0.09 | 0.73 | 0.01 | 27.9 | 7.8 |
| ncRNA_inter_chr15_12600 | 0 | 0.37 | 0 | 0 | 0.02 | 0 | 0.03 | 0.37 | 0 | 0.0 | 11.8 |
| ncRNA_inter_chr15_12601 | 0.01 | 0.31 | 0 | 0 | 0 | 0.06 | 0.00 | 0.31 | 0 | inf | inf |
| ncRNA_inter_chr15_12602 | 0 | 0.34 | 0 | 0 | 0.01 | 0.03 | 0.02 | 0.34 | 0 | 2.6 | 15.1 |
| A1bg | 4.35 | 1,422.30 | 846.40 | 1,422.30 | 1,238.83 | 382.97 | 250.99 | 37.52 | 522.16 | 0.31 | 0.15 |

Highlighting = relative within column for lncRNAs only (excluding A1bg)
NE = (Female Nuclear FPKM / Female Total FPKM)

#### F. FC of Significant Sex Bias (log2; M/F)

| | Log2 FC (M/F) from EdgeR: PolyA Unstr F (G83) | PolyA Unstr F (G85) | Total PolyA Strnd F | Nuclear PolyA Strnd F | Total RiboM Strnd F | Nuclear RiboM Strnd F | B6 Total RiboM Strnd F | # of datasets sign. in |
|---|---|---|---|---|---|---|---|---|
| ncRNA_inter_chr15_12590 | -6.519 | -6.870 | -8.203 | -10.232 | -7.557 | -9.850 | -4.453 | 7 |
| ncRNA_inter_chr15_12591 | -4.937 | -8.111 | -5.436 | -8.313 | -6.344 | -8.715 | -4.078 | 7 |
| ncRNA_inter_chr15_12592 | -6.358 | -6.359 | ns | -5.682 | -5.331 | -7.359 | ns | 5 |
| ncRNA_inter_chr15_12593 | ns | ns | ns | -7.893 | -5.347 | -8.189 | -3.330 | 4 |
| ncRNA_inter_chr15_12595 | ns | ns | ns | -6.804 | -5.586 | -6.170 | ns | 3 |
| ncRNA_inter_chr15_12596 | ns | ns | ns | -5.284 | ns | -7.703 | ns | 2 |
| ncRNA_inter_chr15_12597 | ns | ns | ns | -6.741 | ns | -7.682 | ns | 2 |
| ncRNA_inter_chr15_12598 | ns | ns | ns | -6.147 | ns | -6.023 | ns | 2 |
| ncRNA_inter_chr15_12599 | ns | ns | ns | -7.073 | ns | -6.459 | ns | 2 |
| ncRNA_inter_chr15_12600 | ns | ns | ns | ns | ns | -5.199 | ns | 1 |
| ncRNA_inter_chr15_12601 | ns | ns | ns | ns | ns | -5.464 | ns | 1 |
| ncRNA_inter_chr15_12602 | ns | ns | ns | ns | ns | -5.388 | ns | 1 |
| A1bg | -7.620 | -10.348 | -8.914 | -9.734 | -7.278 | -7.964 | -12.38 | 7 |

Highlighting = relative within table for sex bias and relative within column for "# of datasets sign. In"
"# of datasets..." = the number of RNA-seq datasets (left) in which significant sex bias is observed (max of 7)

## G.

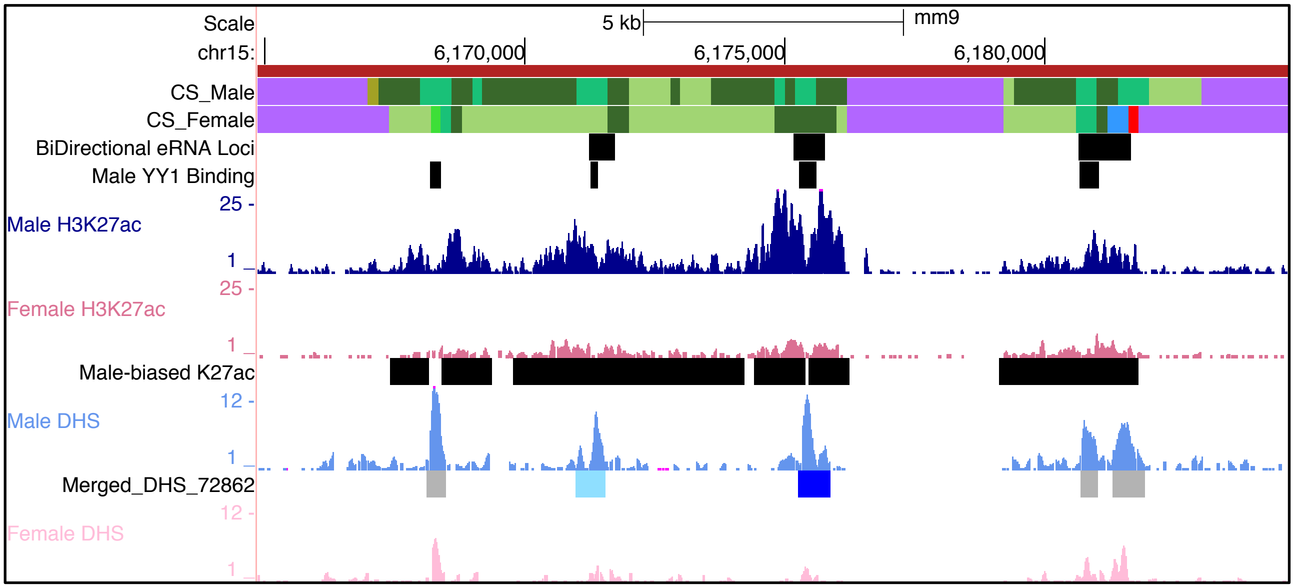

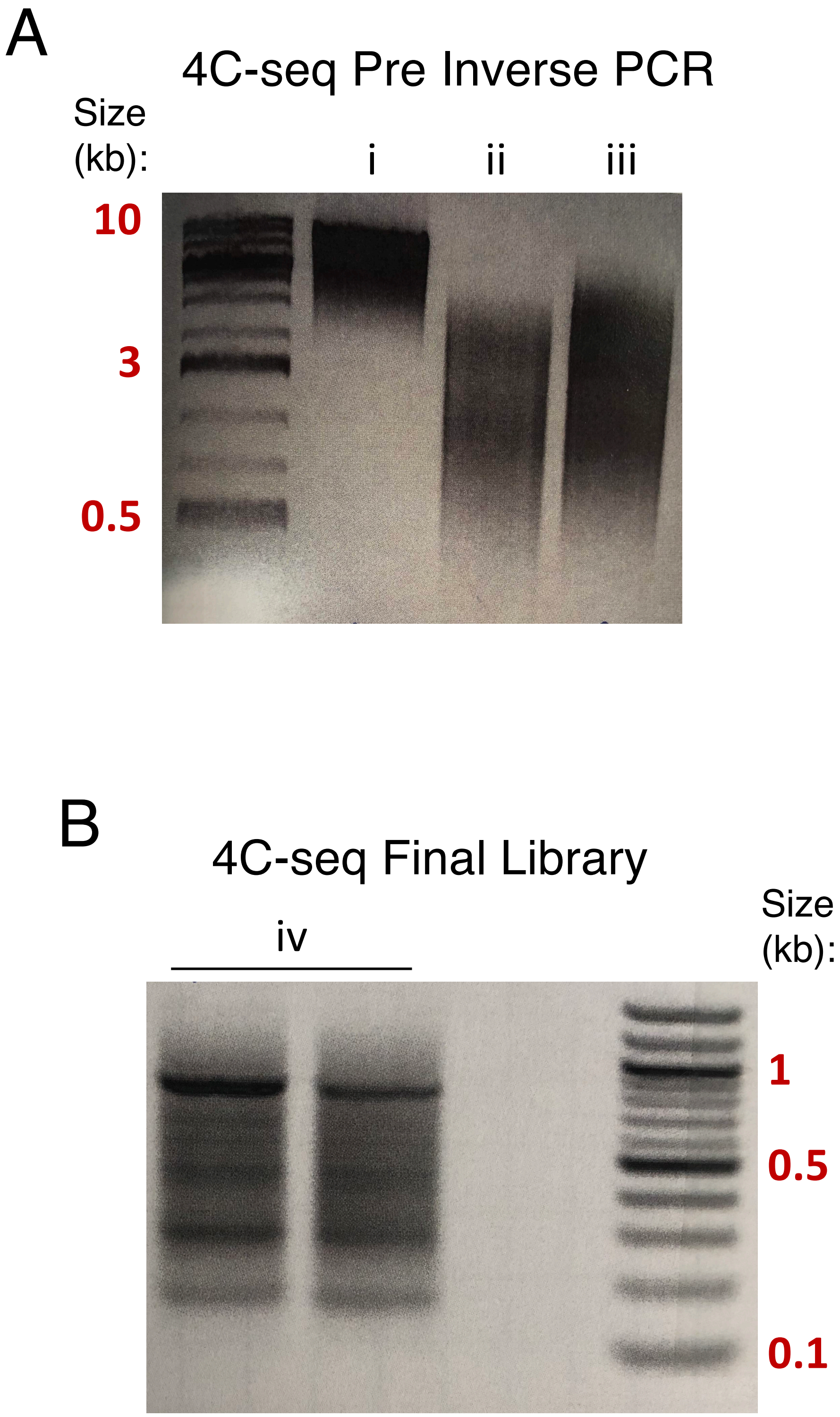
A
4C-seq Pre Inverse PCR
Size (kb):
i
ii
iii
10
3
0.5
B
4C-seq Final Library
iv
Size (kb):
1
0.5
0.1

Fig. S7

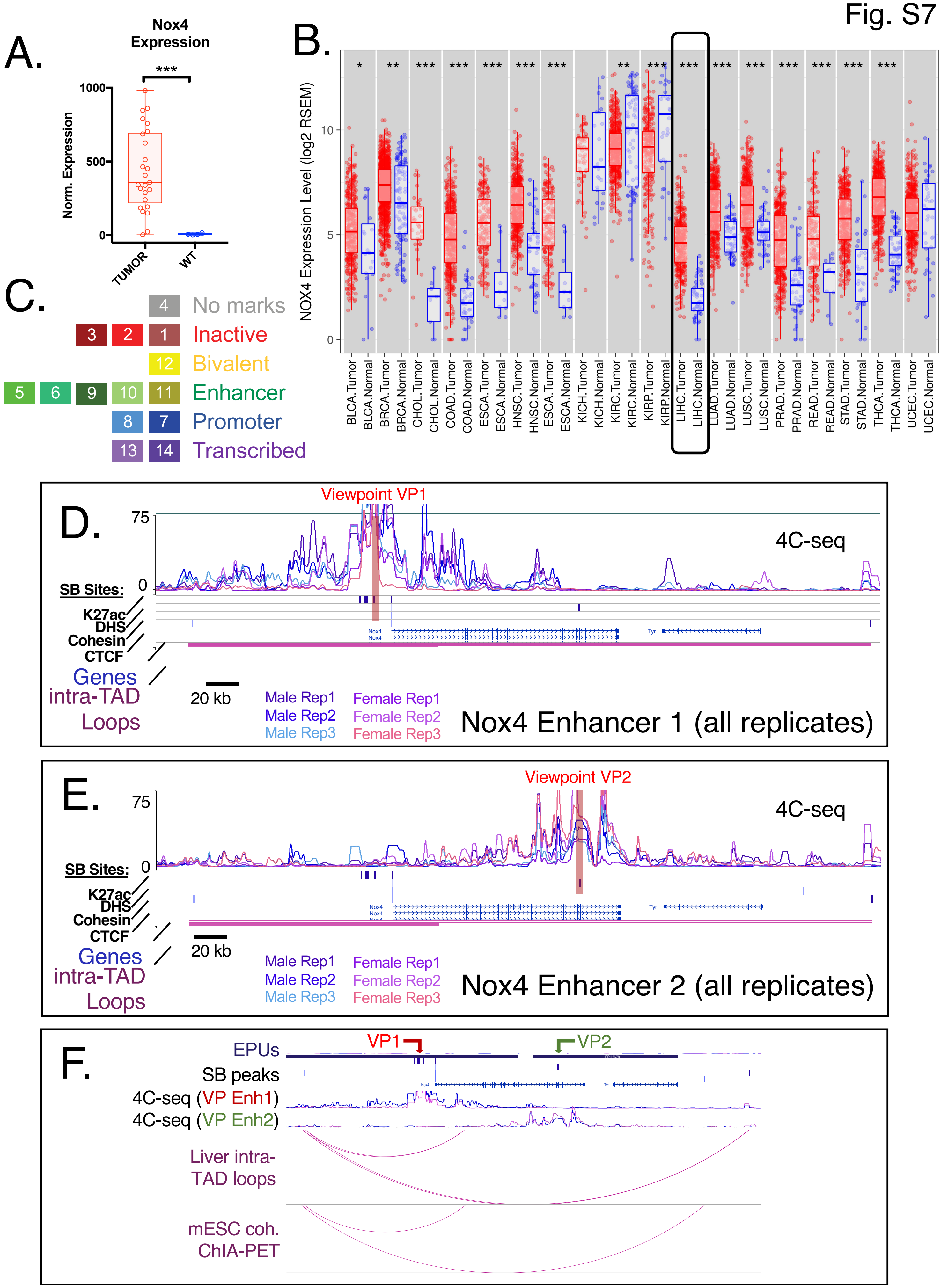

A.
Nox4 Expression
***
Norm. Expression
1000
500
0
TUMOR
WT
B.
NOX4 Expression Level (log2 RSEM)
10
5
0
BLCA.Tumor
BLCA.Normal
BRCA.Tumor
BRCA.Normal
CHOL.Tumor
CHOL.Normal
COAD.Tumor
COAD.Normal
ESCA.Tumor
ESCA.Normal
HNSC.Tumor
HNSC.Normal
ESCA.Tumor
ESCA.Normal
KICH.Tumor
KICH.Normal
KIRC.Tumor
KIRC.Normal
KIRP.Tumor
KIRP.Normal
LIHC.Tumor
LIHC.Normal
LUAD.Tumor
LUAD.Normal
LUSC.Tumor
LUSC.Normal
PRAD.Tumor
PRAD.Normal
READ.Tumor
READ.Normal
STAD.Tumor
STAD.Normal
THCA.Tumor
THCA.Normal
UCEC.Tumor
UCEC.Normal
C.
4 No marks
3 2 1 Inactive
12 Bivalent
5 6 9 10 11 Enhancer
8 7 Promoter
13 14 Transcribed
D.
Viewpoint VP1
75
0
4C-seq
SB Sites:
K27ac
DHS
Cohesin
CTCF
Genes
intra-TAD
Loops
20 kb
Male Rep1
Male Rep2
Male Rep3
Female Rep1
Female Rep2
Female Rep3
Nox4 Enhancer 1 (all replicates)
E.
Viewpoint VP2
75
0
4C-seq
SB Sites:
K27ac
DHS
Cohesin
CTCF
Genes
intra-TAD
Loops
20 kb
Male Rep1
Male Rep2
Male Rep3
Female Rep1
Female Rep2
Female Rep3
Nox4 Enhancer 2 (all replicates)
F.
EPUs
VP1
VP2
SB peaks
4C-seq (VP Enh1)
4C-seq (VP Enh2)
Liver intra-TAD loops
mESC coh. ChIA-PET
